# Robust dimethyl-based multiplex-DIA workflow doubles single-cell proteome depth via a reference channel

**DOI:** 10.1101/2022.12.02.518917

**Authors:** Marvin Thielert, Corazon Ericka Mae Itang, Constantin Ammar, Florian A Schober, Isabell Bludau, Patricia Skowronek, Maria Wahle, Wen-Feng Zeng, Xie-Xuan Zhou, Andreas-David Brunner, Sabrina Richter, Fabian J Theis, Martin Steger, Matthias Mann

## Abstract

Single-cell proteomics aims to characterize biological function and heterogeneity at the level of proteins in an unbiased manner. It is currently limited in proteomic depth, throughput and robustness, a challenge that we address here by a streamlined multiplexed workflow using data-independent acquisition (mDIA). We demonstrate automated and complete dimethyl labeling of bulk or single-cell samples, without losing proteomic depth. In single runs of mammalian cells, a three-plex analysis of tryptic peptides quantified 7,700 proteins per channel. The Lys-N enzyme enables five-plex quantification at MS1 and MS2 level. Because the multiplex channels are quantitatively isolated from each other, mDIA accommodates a reference channel that does not interfere with the target channels. Our algorithm RefQuant takes advantage of this feature and confidently quantifies close to 4,000 proteins in single cells with excellent reproducibility, while our workflow currently allows routine analysis of 80 single cells per day. The concept of a stable proteome still holds at this deeper proteome coverage.

## Introduction

Characterizing biology directly at the single-cell level is greatly advancing our knowledge of different cell types and cellular heterogeneity. Single-cell RNA sequencing (scRNAseq) is now routine and large atlases of human cell populations of different organs are being generated (Consortium* *et al*, 2022; Eraslan *et al*, 2022; Suo *et al*, 2022). In order to clearly define single cell types or sub-types, such measurements often encompass tens or hundreds of thousands of single-cell transcriptomes.

Single-cell mass spectrometry (MS)-based proteomics (scProteomics) is also generating much interest, because the dynamic proteome is thought to be a very informative reflection of the biological function of cells. Furthermore, scProteomics could complement other omics modalities in a multi-level description of cellular systems. However, this requires overcoming key technological challenges in four areas: lossless sample preparation, high performance chromatography, high sensitivity MS measurements and optimal analysis of the low-level signals for quantification. A pioneering approach called nanoPOTS (nanodroplet processing in one pot for trace samples) coupled sophisticated protein extraction to dedicated, narrow column chromatography (Kelly, 2020). SCoPE-MS employs the tandem mass tag (TMT) isobaric labeling strategy to differentially mass-label the single cells, to which a ‘booster channel’ - originally consisting of hundreds of single-cell equivalents - is added (Budnik *et al*, 2018). By reducing analytical requirements for peptide identification more than hundred-fold, it could readily be employed on standard instrumentation. Quantification is less straightforward in SCoPE-MS because the booster channel also contributes to the reporter ion readout in TMT, and this has been shown to lead to ratio distortions (Cheung *et al*; Ctortecka *et al*, 2022b; Ye *et al*).

Our group recently described an ultra-high sensitivity workflow for single-cell applications that builds entirely upon readily available components (Brunner *et al*, 2022). Samples are deposited into low adsorption 384-well plates, allowing parallel preparation with minimal losses. The resulting peptides are deposited on standard Evotips, from which they are eluted in ‘nano-packages’ of 20 nL into a preformed gradient. This stored gradient is then separated at a very low flow ‘Whisper’ gradient (100 nL/min) on an analytical column attached to the Evosep system (Bache *et al*, 2018) and electrosprayed into a trapped ion mobility time of flight mass spectrometer (timsTOF) (Meier *et al*, 2018). We performed data acquisition by dia-PASEF (Meier *et al*, 2020) and achieved median protein identifications of about 1,000 in interphase HeLa cells and up to 2,000 in drug arrested mitotic cells.

Based on these previous experiences with scProteomics, we aimed to develop an improved workflow building on recent developments in multiplex-DIA (mDIA). Although DIA is overwhelmingly performed in a label-free and single-run manner, researchers have long investigated if some of the advantages of multiplexing could be transferred to the DIA setting as well. The principal challenge in mDIA is that multiplexing further multiplies the complexity of already highly complex DIA spectra. To our knowledge the MEDUSA method first reported this concept. Here, ubiquitin and SUMO peptides were differentially labeled with mTRAQ reagents (Δ0, Δ4, and Δ8) (Griffiths *et al*, 2014). Garcia and colleagues combined DIA with SILAC labeling achieving similar identification depth with an order of magnitude better quantitative accuracy than SILAC alone (Pino *et al*, 2021). Very recently, a powerful triplex mTRAQ workflow has been described, which approached similar proteome depth as label-free DIA data (Derks *et al*, 2022) and exceeded its quantitative precision within runs. Applied to single cells, their ‘plexDIA’ approach reached a depth of 1,000 proteins per cell. Apart from these non-isobaric multiplexing methods, low mass reporter based isobaric methods or precursor coupled reporter tags have also been described (Ctortecka *et al*, 2022a; Tian *et al*, 2020).

Here we present a mDIA workflow that uses dimethyl labeling for multiplexing. This derivatization has been well established in proteomics from the original report (Hsu *et al*, 2003) and extensive subsequent work by the Heck group (Boersema *et al*, 2008, 2009; Taouatas *et al*, 2010). We describe the principles of dimethyl mDIA, characterize its performance at the bulk proteomics level on modern MS instrumentation and extend to five-plex at the MS1 and MS2 level. Next, we develop an automated workflow using robotic derivatization combined with Evotips for single-cell proteomics. We then evaluate the concept of a reference channel in mDIA – in which one of the channels is used as a spike-in proteome, making measurements universally comparable to each other. In the context of single-cell proteomics it doubles identifications and throughput. Finally, we devise Reference Quantification (RefQuant) an algorithmic strategy to make optimal use of the reference channel for quantification and apply it to single-cell proteomics.

## Results

### Exploration of dimethyl labeling-based multiplexed data-independent acquisition (mDIA)

Previous approaches to multiplexed data-independent acquisition (mDIA) mass spectrometry have used SILAC or amino-reactive labels, specifically the non-isobaric analog to the iTRAQ reagent, termed mTRAQ (Griffiths *et al*, 2014; Derks *et al*, 2022). To extend the repertoire of available tools for multiplexed analysis of complex proteomes by DIA-MS, we investigated whether a complementary chemical labeling approach based on peptide dimethylation was also suited for this. We decided to explore dimethylation as the derivatization of peptides with dimethyl labels, which is quick, reliable, cost-efficient, and can easily be automated (Raaijmakers *et al*, 2008). Primary amines occurring on peptide N-termini or epsilon-amino groups of lysine residues are thereby derivatized through formaldehyde, to form an intermediate Schiff base that is subsequently reduced by sodium cyanoborohydride, to form a dimethylamino group (Hsu *et al*, 2003). Depending on the combination of stable isotope labeling reagents, this adds a 28.0313 Da, 32.0564 Da, or 36.0757 Da mass tag to each amino group in a given tryptic peptide (referred to as light (Δ0), intermediate (Δ4) and heavy labels (Δ8)), enabling three-plexed mDIA of proteomes (Fig 1A).

**Fig 1:**
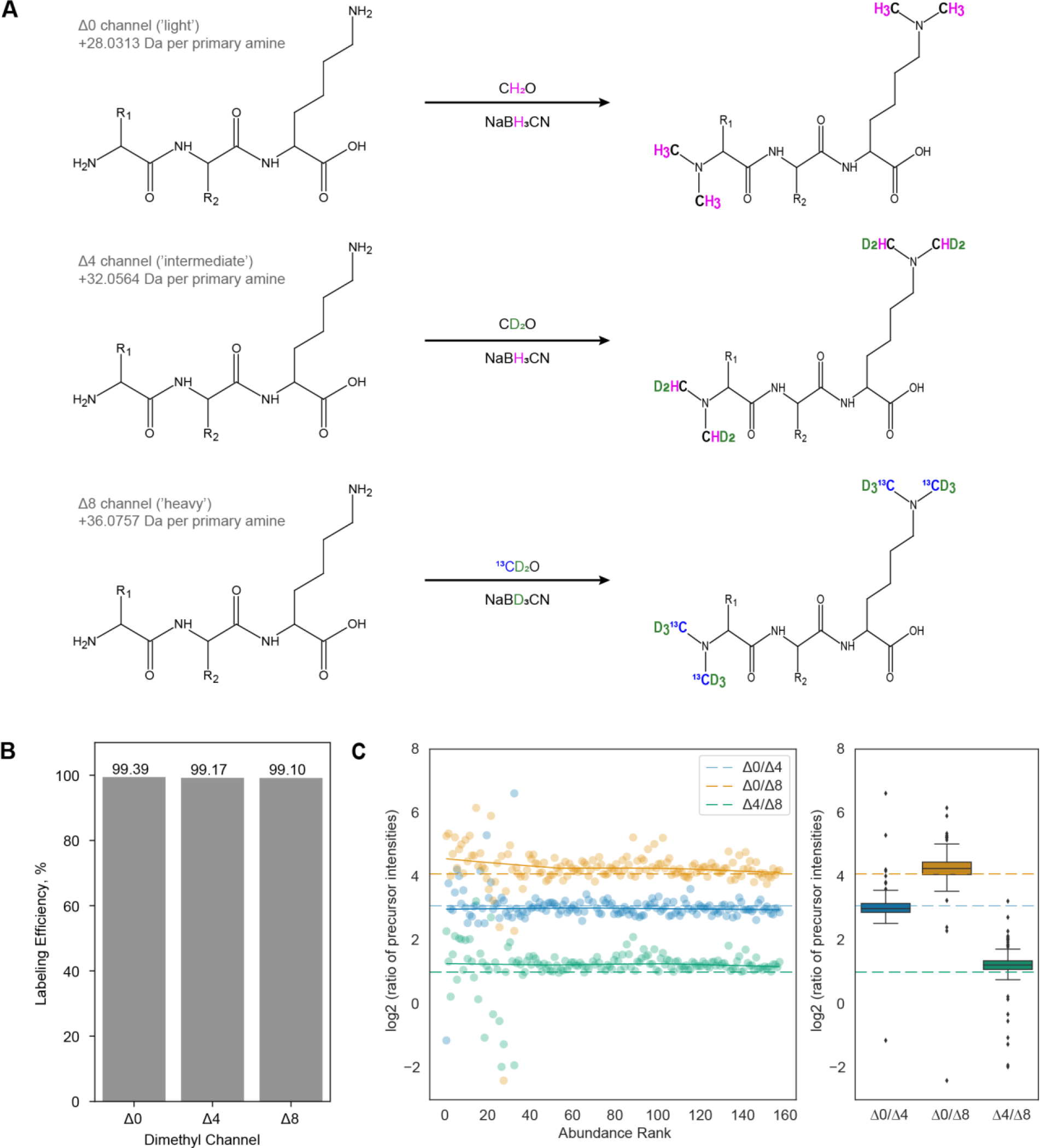
Dimethyl labeling of bovine serum albumin (BSA) combined with data-independent acquisition (mDIA). A. Stable isotope dimethyl labeling scheme for a three-plex mDIA setup. Six hydrogens can be replaced by deuterium and the two carbons by 13C per dimethyl, of which there are one in tryptic peptides ending in arginine and two in those ending in lysine. Depending on the combination of stable isotope labeling reagents, mass tags of 28.0313 Da, 32.0564 Da, or 36.0757 Da are added to each primary amine group of a peptide. A tryptic peptide harboring a C-terminal lysine residue is depicted. B. Dimethyl labeling efficiency of peptides derived from intensity ratios of labeled peptides relative to all detected peptides in DDA mode. C. Quantification accuracy of dimethyl labeled peptides in DIA mode. Differentially labeled tryptic BSA peptides were mixed in a 17:2:1 ratio of Δ0/Δ4/Δ8 and the data was acquired in DIA mode. Scatterplots on the left illustrate the log2 intensity ratios as a function of the peptide abundance rank. In both panels, the expected ratios are marked by colored dashed lines.

Following a previously established dimethyl labeling protocol (Boersema *et al*, 2009), we first evaluated the feasibility of combining it with mDIA. As a first step, we labeled tryptic peptides derived from bovine serum albumin (BSA) with dimethyl-Δ0/Δ4/Δ8, which resulted in a labeling efficiency of more than 99% (Fig 1B). To determine quantification accuracy in this setup, we prepared three samples combining the three channels in different ratios (Δ0/Δ4/Δ8 = 17:2:1; 7:2:1 and 5:3:2) and acquired the data in DIA mode (See Material and Methods), followed by raw data processing with DIA-NN, a neural network-based software that recently has been benchmarked for multiplexed DIA-MS (Derks *et al*, 2022; Demichev *et al*, 2019, 2022). This revealed excellent quantification accuracy, even for the highest tested ratio (17:1) (Fig 1C) and as expected deviations from the true ratios were mainly observed for low-abundant peptides (Fig EV1). We therefore concluded that dimethyl labeling might also be suited for multiplexed acquisitions of complex proteomes in DIA mode.

### mDIA applied to in-depth quantification of complex proteomes

To determine the suitability of dimethyl labeling for mDIA of whole cell protein extracts, we derivatized tryptic peptides from HeLa cells with three dimethyl mass tags (Δ0, Δ4 and Δ8). Like the labeling of just one protein (Fig 1B), this resulted in an almost complete labeling (>99%) of all detected peptides for all three channels (Fig EV2).

Next, we assessed whether identification rates and quantification precision might be compromised by dimethyl labels compared to label-free analysis. First, we measured triplicates of unlabeled and dimethyl labeled peptides (light channel only, Δ0) and processed the raw data with DIA-NN using spectral libraries predicted by AlphaPeptDeep (Zeng *et al*, 2022). Those predicted libraries account for potential differences in peptide fragmentation, collisional cross sections, and retention times between unlabeled and dimethyl labeled peptides.

We used a 75 minute LC gradient and data acquisition either by regular DIA on an Orbitrap platform, or using a timsTOF HT instrument, with our recently reported optimal dia-PASEF acquisition schemes (Skowronek *et al*, 2022). Whereas the timsTOF platform generally yielded higher identification numbers and greater quantification precision, the number of quantified precursors and protein groups between unlabeled and one-channel dimethyl labeled samples was similar on both instruments. This demonstrates that derivatization of peptides with dimethyl groups does not negatively impact peptide identification rates (Fig 2A-C). Using the Orbitrap instrument, we quantified about 7,000 protein groups (85,000 precursors) from 125 ng of injected peptides in both unlabeled and dimethyl-Δ0-labeled samples, with a median CV of 4.8%. In contrast, the timsTOF yielded about 20% more protein groups and 50% more precursors in unlabeled samples (about 8,400 protein groups and 128,000 precursors) and 11% more protein groups and 40% more precursors (7,700 protein groups and 118,000 precursors) in labeled samples with the same injection amount (Fig 2A-B).

**Fig 2:**
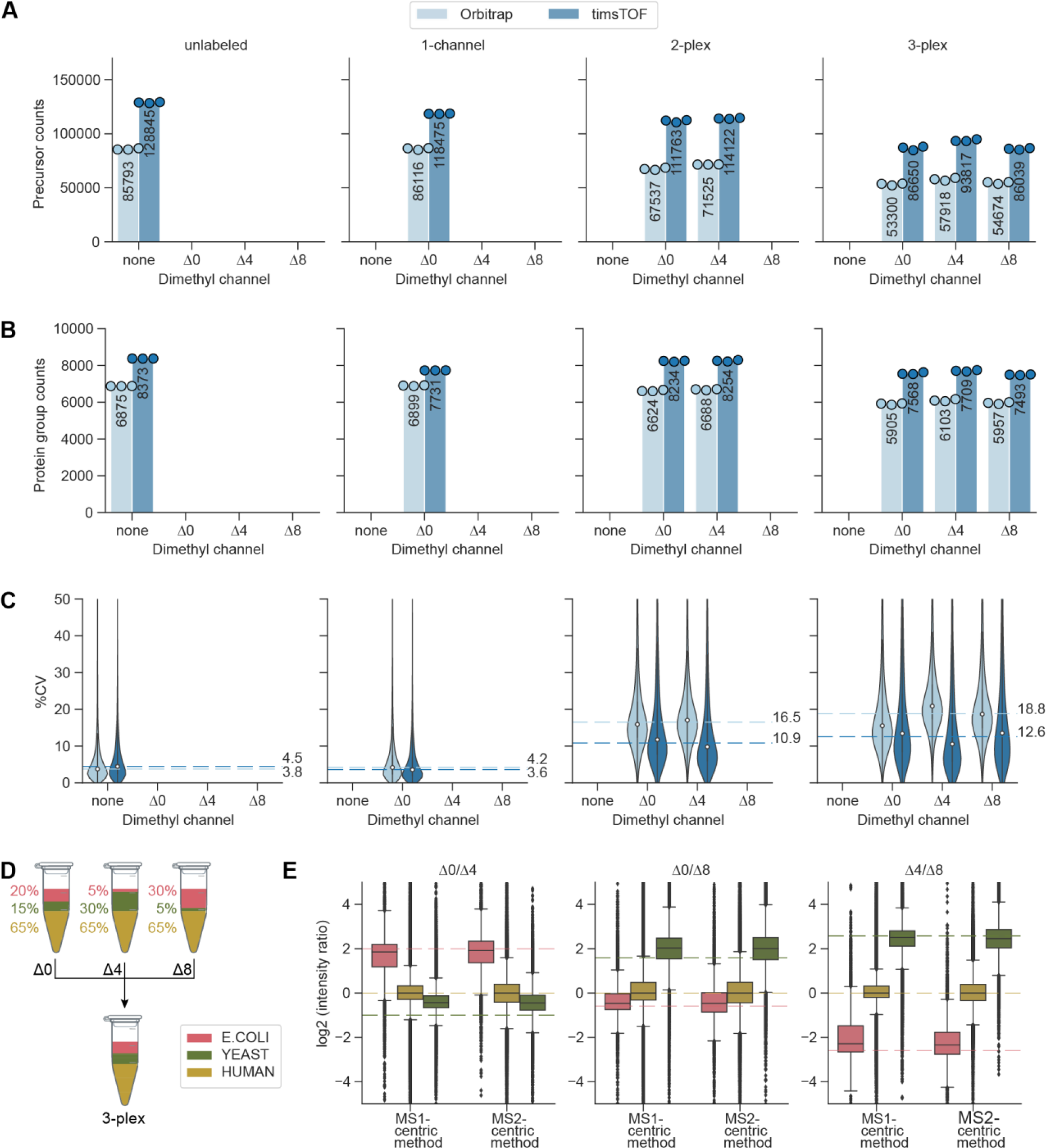
Identification rates, quantification precision and accuracy of dimethyl labeled peptides. A and B. Number of quantified HeLa peptide precursors (A) and protein groups (B) for unlabeled, one-channel (Δ0), two-plex (Δ0 and Δ4), and three-plex (Δ0, Δ4, and Δ8) labeled samples. 125 ng of peptides were injected per channel in technical replicates. C. Coefficients of variation (CV, %) of protein groups shown in (B). Median CVs are shown as dashed lines. D. Mixing scheme for tryptic peptides of HeLa, *S. cerevisiae*, and *E. coli* at different ratios prior to dimethyl labeling. The three channels were multiplexed in a 1:1:1 ratio and 300 ng total amount was measured in triplicates on the timsTOF platform using MS1- and MS2-centric methods, as depicted in Fig EV4. E. Side-by-side comparison of quantification accuracies between MS1-centric and MS2-centric acquisition methods in the mixed species experiment. Protein group ratios are plotted as boxplots.

When increasing proteome complexity by adding a second channel (Δ4), identifications numbers remained almost constant at both the precursor and the protein level on the timsTOF platform (about 115,000 precursors and 8,200 protein groups), whereas we observed a slight decrease on the Orbitrap instrument (about 20% for precursors and 4% for proteins groups). In a three-plex mDIA setup, protein group identifications further decreased on the Orbitrap and slightly decreased with the timsTOF (Fig 2B), indicating that the deconvolution of multiplexed spectra is challenging and that the ion mobility dimension of the timsTOF platform helps to resolve this complexity.

To investigate quantification accuracy in a three-plex setup we combined tryptic peptides from human (HeLa), *S. cerevisiae*, and *E. coli* at defined ratios for three labels (Figure 2D). This mixing scheme creates a benchmark of known protein ratios, which can be compared with the ratios of measured intensity values. In addition to our initially designed dia-PASEF method, we generated an alternative method, consisting of multiple MS1 scans in between dia-PASEF scans of each duty cycle (Fig EV4). We used this MS1-centric method for direct comparison with our standard MS2-centric method, by quantifying protein ratios across channels in the three-plex, mixed species experiment. This revealed that with both methods the measured protein group ratios largely agreed with the expected ones, and that the MS2-centric method slightly outperformed the MS1-centric method in terms of accuracy, with similar identification rates (Fig 2E and Fig EV3). However, MS1-based quantification (MS1-centric method) outperformed MS2 level quantification (MS2-centric method) in quantification precision, making it an attractive option in multiplexed DIA experiments (Fig EV3).

### Extending the scope of multiplexing by dimethyl labels

Although trypsin is the most frequently used protease for MS-based proteomics applications, in the context of dimethyl labeling it limits multiplexing to only three channels if a mass difference between channels of 4 Da is desired at MS1 and MS2 level. At the MS1 level alone, Lys-C in combination with different isotopes of formaldehyde and cyanoborohydride enables five-plex dimethyl labeling (Wu *et al*, 2014).

Lys-C derived peptides harbor two primary amino groups for labeling - one of them is located on the N-terminus and the second one on the side chain of the C-terminal lysine residue. Digesting proteins with Lys-N instead generates a N-terminal lysine on each peptide, and thus two primary N-terminal amines for labeling (Raijmakers *et al*, 2010). Five-plex labeling of Lys-C or Lys-N-derived HeLa peptides thus results in isotopologues that are separated by 4 Da from each other, which would be sufficient for accurate quantification (Fig 3A).

**Fig 3:**
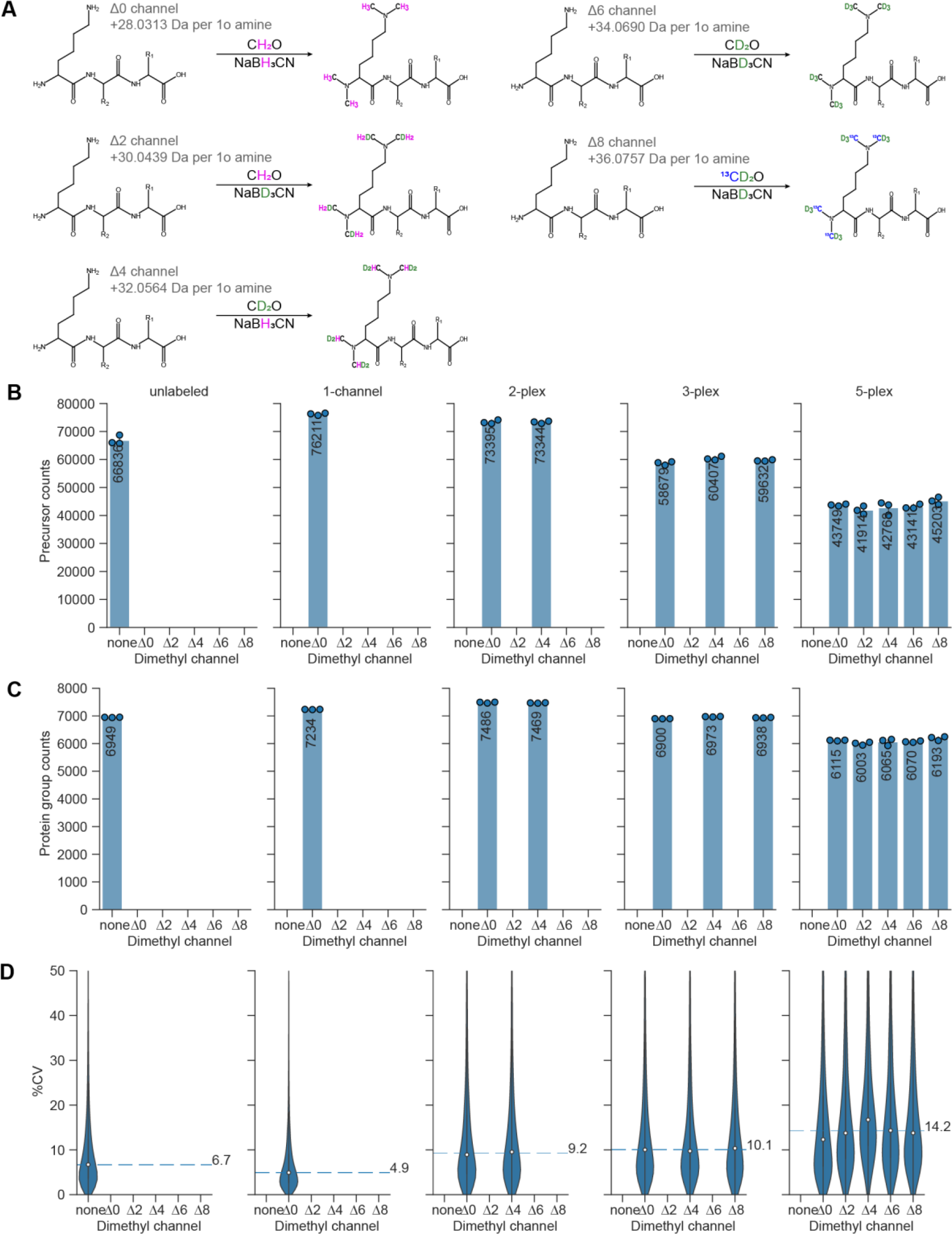
Five-plex dimethyl labeling using Lys-N protease. A. Stable isotope dimethyl labeling scheme for a five-plex mDIA experimental setup with Lys-N digested peptides. Depending on the combination of stable isotope labeling reagents, mass tags of 28.0313 Da, 30.0439 Da, 32.0564 Da, 34.0690 Da, or 36.0757 Da are added to each primary amine group on a peptide. Since Lys-N hydrolyses the N-terminal side of lysine residues, two labels are clustered on the N-termini of peptides. B and C. Number of quantified HeLa peptide precursors (B) and protein groups (C) for unlabeled, one-channel (Δ0), two-plex (Δ0 and Δ4), three-plex (Δ0, Δ4 and Δ8), and five-plex (Δ0, Δ2, Δ4, Δ6 and Δ8) labeled samples. 100 ng of peptides were injected per channel. D. Coefficients of variation (CV, %) of protein groups shown in (C). Median CVs are shown as dashed lines.

Lys-N-derived and dimethylated peptides form more b-ions during MS2 fragmentation than tryptic peptides (Taouatas *et al*, 2008) and the two dimethyl labels are carried by b-ions. We reasoned that this should enable MS2-based quantification in five-plex experiments. In contrast, fragmented five-plex labeled peptides derived from Lys-C carry one dimethyl label on each b- and y-ion. Each of them separates from its isotopologues by only 2 Da, which leads to major overlaps of isotope clusters.

We therefore explored whether Lys-N-mediated digestion combined with five-plex dimethyl labeling could be a straightforward way of further increasing multiplexing for DIA-MS. We used HeLa lysates and digested them with Lys-N, followed by derivatization with five-plex dimethyl mass tags, achieving more than 97% labeling efficiency (Fig EV5). After combining the labeled peptides in equal ratios, we quantified them using the timsTOF instrument, with dia-PASEF as scan mode and employing methods optimized for Lys-N digestion with py_diAID (Fig EV6A) (Skowronek *et al*, 2022). This led to the quantification of more than 7,000 protein groups in both unlabeled and labeled samples with just one-channel (Δ0), and median CVs below 10% (Fig 3B-D). In contrast to dimethyl labeled tryptic peptides, combining up to three labels did not significantly decrease protein group identifications and even in the five-plex setup, protein identifications decreased by only about 15% compared to one-channel dimethyl measurements. This demonstrates the feasibility of multiplexing up to five samples in mDIA experiments using dimethyl labeling.

### Robust and automated ultra-high sensitivity mDIA workflow with a reference channel

Having established the dimethyl mDIA derivatization for bulk proteomics samples, we set out to combine it with our recently published ultra-high sensitivity single-cell proteomics workflow (Brunner *et al*, 2022). We used a 384-well format for all ultra-high sensitivity experiments to reduce reaction volumes. Furthermore, we adapted the lysis and digestion buffer to an amine-free buffer using triethylammonium bicarbonate (TEAB) to enable the amine directed chemistry of dimethyl labeling (Fig 4A). The dimethylation reaction itself is performed by simple addition of the chemicals to the peptide mixtures in small volumes achieving a labeling efficiency greater than 99.5% for all channels at 1 ng tryptic HeLa peptides (Fig 4B). Subsequent cleanup of the derivatization reaction has previously been done in an in-column format (Boersema *et al*, 2009), but such a step is generally omitted in single-cell applications to avoid sample loss. In our format, however, the cleanup and combination of the separately labeled single cells can easily be performed in the Evotips, on which peptides are anyway deposited. Thus, the derivatization reaction does not complicate the workflow. The entire sample preparation (Fig 4A), including loading of the Evotips, was automated on a standard Bravo robot (Agilent) to streamline the workflow, enhance reproducibility and automate for increasing throughput. This also adds traceability and enables compliance and quality control.

**Figure 4:**
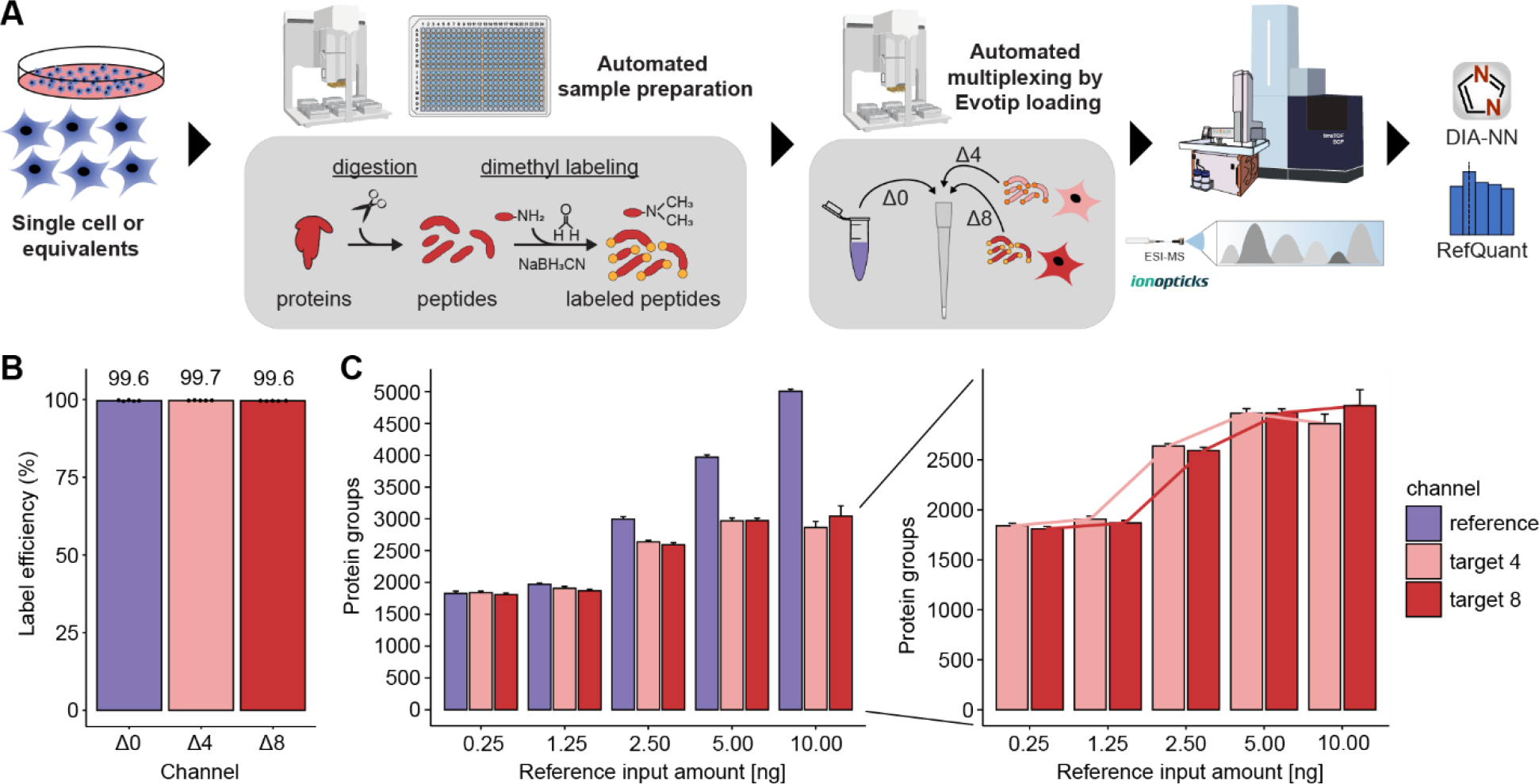
Streamlined and automated ultra-high sensitivity mDIA workflow. A. Single HeLa cells or single-cell equivalents are processed in a standard 384-well plate on a Bravo robot (Agilent) by lysis in TEAB and ACN, tryptic digestion, dimethyl labeling and multiplexing by loading the different mDIA channels onto the same Evotips. LC-MS analysis is done by an Evosep One with low-flow chromatography coupled to a timsTOF SCP instrument. The data is analyzed by DIA-NN, followed by our algorithm RefQuant (see main text). B. 1 ng tryptic peptides from HeLa cells were labeled with dimethyl mass tags Δ0, Δ4 and Δ8 and acquired individually in DDA mode to determine labeling efficiency. Efficiency is calculated based on intensity ratios of labeled peptides relative to all detected peptides. The labeling efficiencies were consistently higher than 99.5 % for all channels in quintuplicate measurements. C. Effect of varying the protein input in the reference channel for protein identification across all channels. Increasing the total protein amount in the reference channel linearly increases protein identifications in the reference channel (left), but importantly protein identifications reach a plateau in the target channels (Δ4 and Δ8) with single-cell equivalents upon 5 to 10 ng. Connected lines between increasing reference input amounts show a sigmoidal relation (right). Error bars represent the standard deviation of quintuplicate measurements.

In the Evosep instrument, peptides are eluted from the tips in nano-packages and analyzed at a very low ‘Whisper’ flow (100 nL/min) with a gradient of 31 min and overhead time to the next injection of 7 min (‘40 samples per day method’). With such a low flow rate, it is important to avoid any post-column dead volumes. We therefore sought to eliminate the post-column connectors that we had employed previously by using pulled columns packed into the electrospray tip (IonOpticks) (Sandow *et al*, 2021). This setup makes the workflow very robust and reproducible, and also improved chromatographic resolution, with peaks eluting in full width half maximum (FWHM) with a median of about 2 s, corresponding to an elution volume of only 10 nL.

Finally, we analyzed the multiplexed single-cell samples using an optimal dia-PASEF method (Skowronek *et al*, 2022) with eight dia-PASEF scans and a mass range of 300 to 1200 and ion mobility of 0.7 to 1.3 on the timsTOF SCP.

In the bulk experiments above, we had established that the different non-isobaric channels in the mDIA workflow are decoupled from each other in terms of quantification. We therefore reasoned that we could use one of the mDIA channels as a ‘reference channel’ comprised of the same input material across all samples. Apart from providing a common standard for identification and quantification, such a reference channel conceptually decouples identification from quantification. This is because the reference channel proteome is easily identified due to its higher signal intensity and its uniformity across samples. Subsequently, the software needs to transfer the quantification boundaries from the reference channel to the target channels containing the single-cell proteomes.

To investigate the reference channel concept in mDIA, we first systematically increased its loading in the Δ0 channel. With our current chromatographic set-up, the number of identifications leveled off at roughly 10 ng and about 5,000 proteins, which we therefore chose as the reference channel amount for all subsequent experiments (Fig EV7A).

Next, we kept the reference amount constant, but varied the amount of peptide in the target channels to explore how much the reference channels would support identifications of weak signals. The DIA-NN software that we used for analysis does not directly have a notion of a reference channel, but considers the boundaries of the channel with the highest scoring precursor as internal reference for transfer. To assess identification confidence, DIA-NN reports a ‘Channel.Q.Value’ parameter (Fig EV7). We experimentally determined the value of this parameter at which features from the reference channel could safely be transferred into the target channels. To this end, we measured mDIA samples in which we left one or both of the target channels empty. This revealed that a ‘Channel.Q.Value’ of 0.45 led to a count-based FDR of 1% for precursors and 0.17 for protein groups (Suppl. Fig. EV7C). At the latter value, we also obtained accurate quantitative ratios between the target and reference channel as judged by defined mixing experiments (Fig EV7D). To ensure high confidence identifications in the target channels with single-cell equivalents and good quantification, we henceforth filtered the data with a ‘Channel.Q.Value’ of 0.15 as well as the recommended parameters of DIA-NN.

With these parameters, we investigated whether increasing amounts in the reference channel might influence the number of proteins found in single-cell equivalents in the target channels. Importantly, while increasing the reference channel input amount beyond 5 ng the target channel identifications remained the same. (Fig 4C). We conclude that these channels are isolated from each other as expected from the mDIA concept.

### RefQuant improves quantification by sampling ratios relative to the reference channel

Having shown the advantages of a reference channel to substantially increase protein identifications in the target channels (Fig. 4C), we investigated if its benefits also extend to quantification. We reasoned that the standardized reference proteome present at higher amounts might directly improve quantification in the target channels by effectively reducing technical variation across runs – an idea already implemented in the Super SILAC strategy (Geiger *et al*, 2010). To enable this analysis, we implemented an approach that we term **Ref**erence **Quant**ification (RefQuant), which is based on sampling all available ratios relative to the reference channel (Figure 5A). In short, the ratios of individual fragment ions as well as MS1 peaks between reference and target channels are extracted, which results in a distribution of ratios. Subsequently, a best ratio R is estimated from the ratio distribution in a robust manner by taking the mean from the 40% upper quantile of ratios. We determined this threshold empirically using the scBenchmark dataset (see below) (Fig EV8B). The intensity of the target channel is then calculated by multiplying R with a scaling factor representing the intensity of the reference channel. This gives a re-scaled precursor intensity value. The scaling factor is derived per precursors as the median reference intensity over all runs, in order to further stabilize the estimate. An interesting property of RefQuant is that it utilizes the high relative ratio to the reference channel in two ways: 1) the prior knowledge about the high expected ratio is used as an effective ‘noise filter’ (Fig EV8A-D) and 2) the higher abundance of the reference channel stabilizes quantification, which should conceptually reduce overall noise (Text EV9).

**Figure 5:**
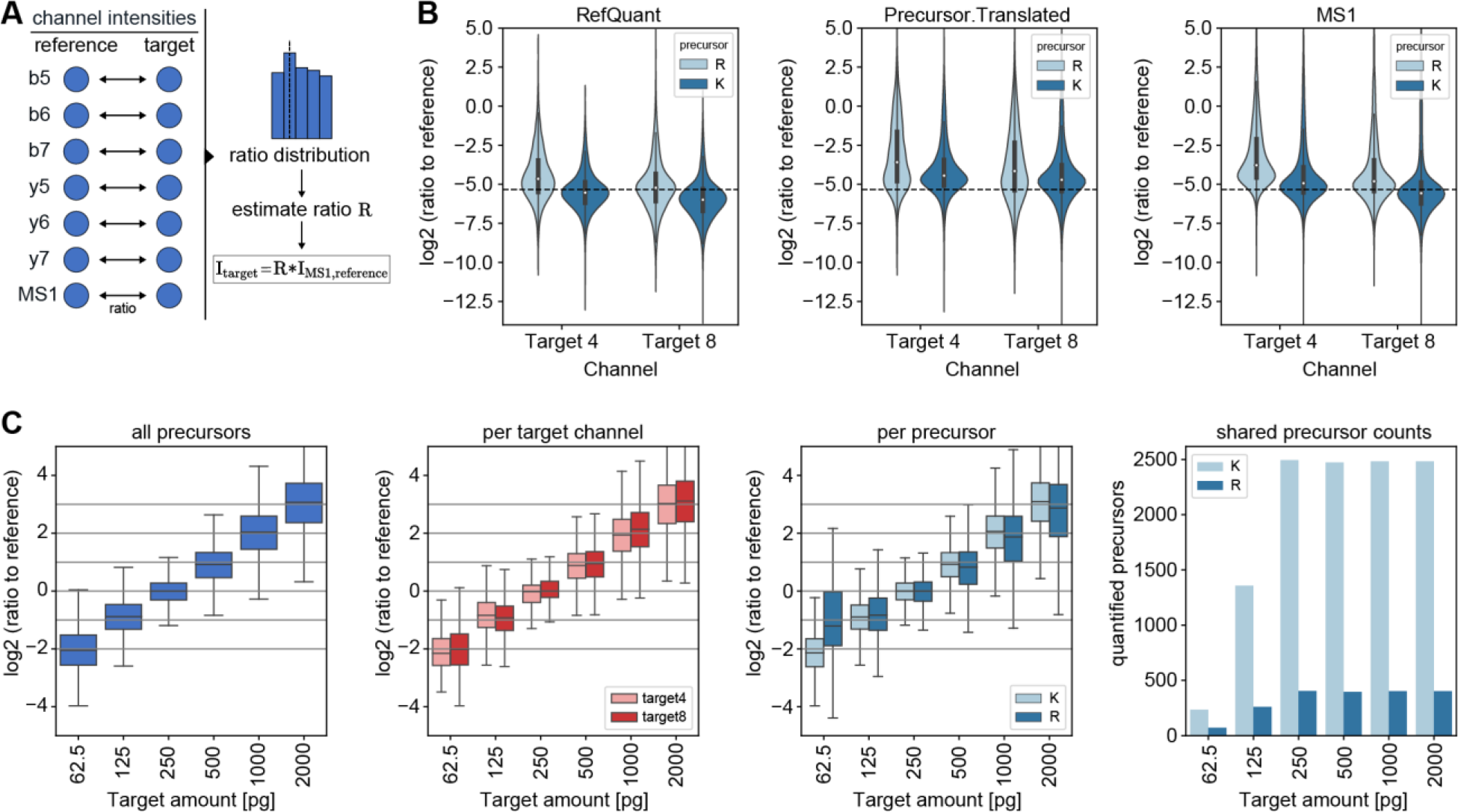
RefQuant quantifies single-cell equivalents accurately based on sampling ratios relative to the reference channel. A. Concept of RefQuant in calculating ratios between each fragment and MS1 peak for each precursor between reference and each target channel. The resulting ratio distribution is filtered for the first 40% of the quantile and the ratio factor R is calculated by the mean of the resulting distribution. This ratio factor R is then multiplied by the reference intensity to retrieve the RefQuant target intensity. B. Ratio of reference channel to each target channel by arginine and lysine precursor based on RefQuant (left), Precursor.Translated by DIA-NN (middle) and MS1 by DIA-NN (right). RefQuant showed best the expected ratios in both target channels compared to DIA-NN quantities of Precursor.Translated and MS1. C. RefQuant accurately quantified four-fold differences in protein amount between reference and target channels (scQuant dataset) and correctly extracted the expected ratios across channels for lysine and arginine precursors. The ratio of reference to target channel was calculated using RefQuant. All log2 ratios were normalized to the single-cell equivalent sample in the target channel (250 pg).

To test RefQuant, we initially applied it to a single-cell equivalent benchmark dataset (scBenchmark, target channels with 250 pg HeLa peptides) in each of the target channels and one in which we varied the amount of HeLa peptides covering four-fold expression differences in both directions from a single-cell equivalent (scQuant, target channels from 62.5 to 2000 pg HeLa peptides). Note that the summed protein intensity per run in the scBenchmark and scQuant dataset is lower than the intensity in our single-cell dataset (see next section), which indicates that the true mass of our single HeLa cells is indeed close to 250 pg or even slightly larger, making our scBenchmark and scQuant at least as challenging to measure as the true single cells.

In the scBenchmark dataset, the ratio between the reference and target channel is well reconstructed by RefQuant, as compared to the standard method of only extracting the target ion intensities without using the reference channel (Fig 5B). This indicates good reproducibility and also good quantitative accuracy. This was the case both for MS1 (Ms1.Area) and MS2 (Precursor.Translated) quantification. In particular, quantification without RefQuant had more scatter and RefQuant best reconstructed the true ratios. DIA-NN reports a difference between quantification of tryptic peptides with C-terminal lysine (‘lysine precursors’) and arginine (‘arginine precursors’). RefQuant also accurately quantifies arginine precursors, which are particularly challenging to quantify, presumable because they have shared y-ions between mDIA channels. We next turned to assess the accuracy of different fold changes in single-cell equivalents, the most important outcome of single-cell proteomics. The scQuant dataset shows that RefQuant on mDIA accurately retrieves four-fold differences in the target channel with either decreasing or increasing amount in the target channel. There were no differences in the two target channels (Δ4 and Δ8) after median correction (Fig 5C and Fig EV8E). The quantification of lysine peptides, which have no shared fragments between channels, has less variation compared to arginine precursors although the results of the arginine peptides may be skewed by their small number at 62.5 pg (Fig. 5C).

### mDIA identifies up to 4,000 proteins per single cell in unsynchronized cell populations

Having shown the benefits of the reference channel for identification and quantification, we next applied our mDIA workflow to single cells. We designated one of the channels as the reference (Δ0), leaving two target channels (Δ4 and Δ8) for single-cell analysis. With this setup, we routinely analyzed 80 single cells per day.

The only difference to the ultra-high sensitive mDIA workflow described above is that we upfront sorted single HeLa cells from culture by flow cytometry into 384-well plates. We FACS-isolated the cells in an unbiased manner; thus we expected a range of cell sizes depending on the cell cycle stage. This was reflected in a wide range of protein identifications across analyzed cells (Fig. 6A). To investigate the cause of this spread, we considered the summed MS signal of all peptides as a proxy for total protein amount in our single cells. Plotting this signal against protein identifications revealed a strong dependency of input amount on proteome depth (Fig. 6B). This suggests that further gains in ion signals will allow even deeper coverage in the reference mDIA workflow in the future.

**Figure 6:**
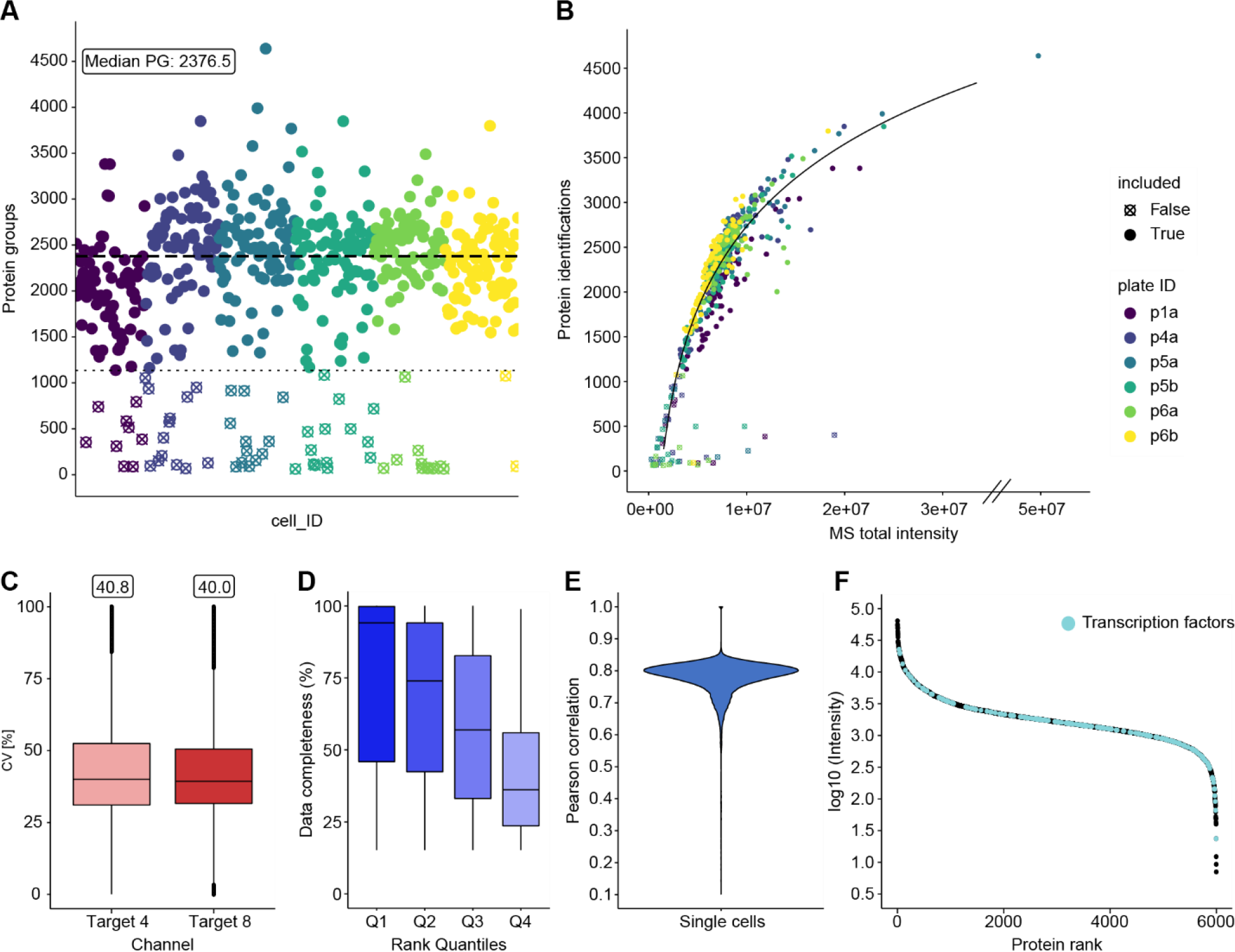
Reference channel enables identification of almost 4,000 protein groups in a single cell. A. Protein identifications of single-cell measurements. 476 single cells reveal a median identification of 2,377 with identifications up to 4,000 protein groups over six 384-well plates (disregarding a single outlier of 4,600 protein groups). B. Protein identifications versus the total sum of MS intensity per cell measurement shows logarithmic dependency of protein identifications upon MS signal intensity. C. CV distribution of single cells in each target channel. D. Completeness of ranked quartiles (Q1: 1-924; Q2: 925-1,848; Q3: 1,849-2,772; Q4: 2,773-3,696). Proteins with high rank show higher data completeness compared to low ranked proteins. E. Pearson correlation of single cells of the mDIA workflow. F. Rank plot of close to 6,000 proteins. Transcription factors are highlighted across abundance range (cyan).

However, even at this stage, we identified almost 4,000 protein groups in single cells (disregarding a single outlier at 4,600 identifications). Across our 476 cells in the entire dataset, we obtained 5,997 proteins, a substantial part of the proteome expressed by a single cell. Even compared to some of the deepest proteomes that were achieved in this cell type with hundreds of µg rather than pg input (Bekker-Jensen *et al*, 2017), our single-cell proteomes together account for half of the proteins.

In median, we reach a proteomic depth of about 2,377 protein groups. This is more than twice the number of the median identified in interphase cells in our previous publication, almost entirely attributable to the improvement due to the reference channel (Fig 6A) (Brunner *et al*, 2022). The overall coefficient of variation due to technical and biological factors was 0.4 (Fig 6D). Overall data completeness was high. After we filtered out proteins that were only detected in 15% of the cells – possibly due to biological variation – data completeness of the remaining 3,696 proteins was 59.3% across the entire dataset. As expected, this depended on the abundance of the proteins, with completeness of the top rank order quartile at 94.1% and the lowest quartile at 36.2% (Fig. 6D). The median overall Pearson correlation of the single cells was 0.79 (Fig 6E).

Overall, our protein signals cover more than four orders of magnitude of dynamic range, in which we identify many transcription factors, like the Hox family, STAT1-3 and SOX6, which range over the whole abundance range, but are mainly expressed at low amounts (Fig 6F). This constitutes to 20% of all human transcription factors (Lambert *et al*, 2018).

### Stable proteome at deeper proteome coverage

In our recent single-cell publication, we discovered evidence for a stable proteome while the transcriptome had a much higher overall variability as measured by CV. We attributed this difference to the fact that single cells need a complete proteome to function, whereas transcripts for genes expressed in these same cells are only needed rarely (Brunner *et al*, 2022). Thus, numbers of transcripts for many expressed genes are often below one on average across many single cells. A limitation of our previous study was the coverage of the single-cell proteome – about 1000 proteins in interphase cells and up to 2000 proteins in drug arrested mitotic cells.

Here, we used the same strategy as in our label-free single-cell dataset (Brunner *et al*, 2022). In brief, we investigated the variability of protein expression by the CV as a function of abundance by MS intensity. Remarkably, we still see relatively small CVs across covered abundances for the whole measured dynamic range (Fig EV10A). Previously, we had defined a ‘core proteome’ of the top 200 proteins with at least 70% data completeness. We also observe lower CVs and higher mean MS intensity for these compared to noncore proteins (Fig EV10B and C). The somewhat increased CVs of the proteins that were only quantified here, is readily explained by their lower signal in the MS.

As before, we next compared our mDIA single-cell dataset to scRNAseq data based on Drop-Seq (Macosko *et al*, 2015) and SMART-Seq2 (Picelli *et al*, 2014) on the same cell type. In the count distribution plot, scRNAseq and mDIA scProteomics data again separate from each other, indicating the stable proteome at higher proteomic coverage (Fig 7A). CV values for the proteins are tight and low while scRNAseq has higher CV values (Fig 7B). Together these data show that the concept of a stable proteome still holds true at the higher proteomic depth achieved here.

**Figure 7:**
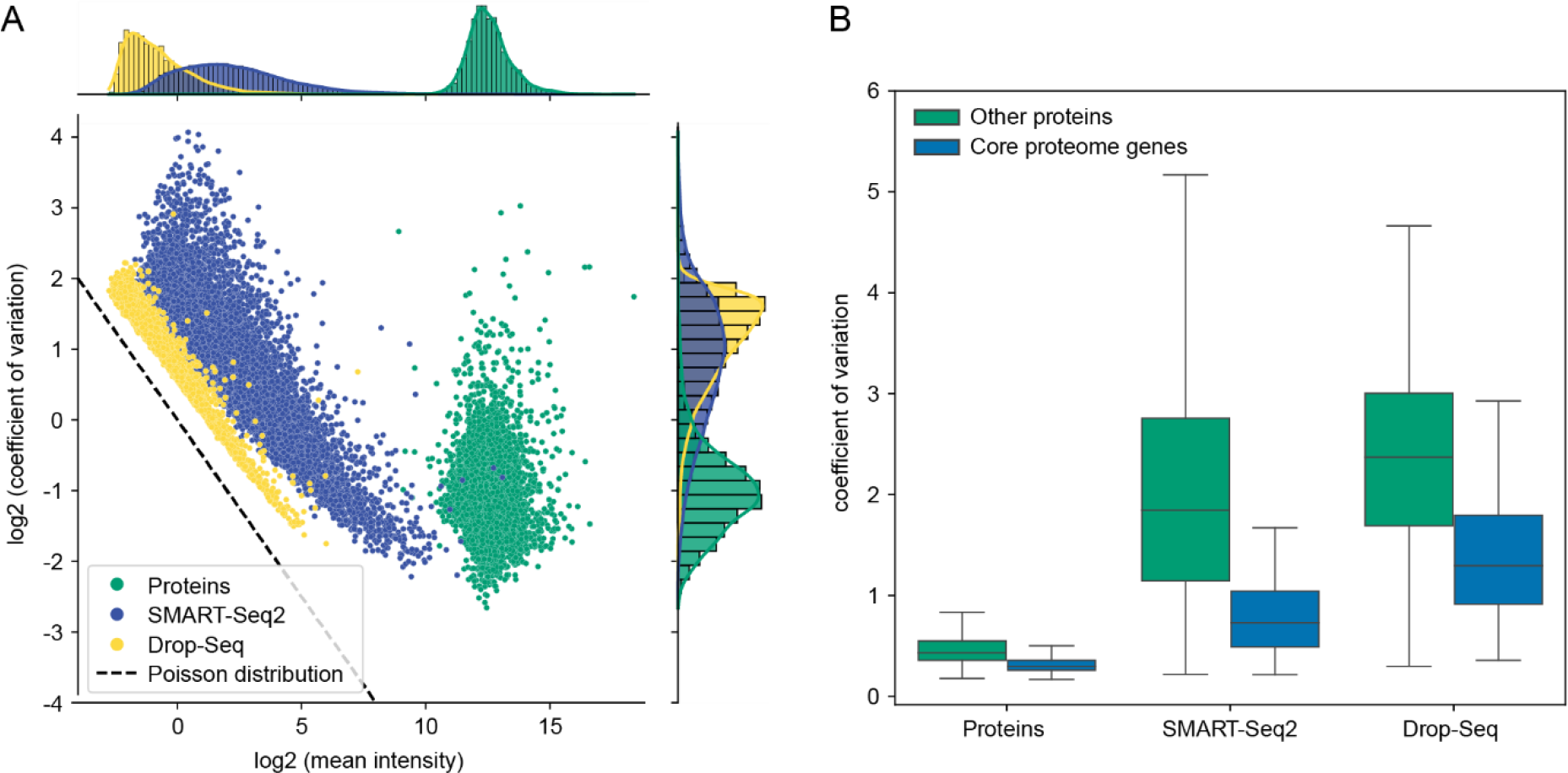
The concept of a stable proteome is still valid at higher proteomic depth in single cells. A. Coefficients of variation of single-cell mDIA protein expression plotted against the mean intensity of each protein for single-cell proteins (green) and scRNAseq of SMART-Seq2 (blue) and Drop-Seq (yellow). B. Comparison of coefficients of variation between single-cell proteomics and scRNAseq with regards to the ‘core proteome’ (blue) and other genes (green).

## Discussion

Here, we described a robust and economical mDIA workflow that we explored using standard proteomics samples and single cells. One of its main advantages is that it uses dimethyl labeling, a well-established, very simple and robust technology (Hsu *et al*, 2003; Boersema *et al*, 2009). We consistently achieved labeling efficiencies of more than 99% for peptides generated by trypsin and Lys-C – proteases typically employed in proteomics. The labeling protocol consists of only three pipetting steps, hence it was easily automatable on a standard liquid handling robot. The differentially labeled peptides are combined into one stage-tip for purification and storage, thus avoiding sample loss. Another practical advantage is that the reagents themselves are very stable, whereas broadly used NHS (N-Hydroxysuccinimide)-chemistry-based reagents such as mTRAQ or TMT are more labile.

Dimethyl labeling enables quantification at both the MS1 and MS2 level, but is limited to three-plex when used in combination with trypsin. Here we show that Lys-N digestion followed by dimethyl labeling accommodates a five-plex format at both levels as well, with the interesting property that b-ions are split up into five quantifiable states separated by 4 Da. The y-ions are less intense due to the absence of a C-terminal lysine, however, this is compensated for identification purposes by the fact that y-ions from all channels stack on top of each other. This will be particularly interesting in the context of fragmentation with Electron Transfer Dissociation (ETD), as this has been shown to generate nearly pure c-ion series in many cases on triply charged precursors (Hennrich *et al*, 2010).

Current software tools do not yet make optimal use of dimethyl mDIA data. Unlike 13C-based labels, dimethyl labels can slightly influence retention times on LC columns. Our results show that this is not a major impediment to quantification, but we believe that results will improve once the retention times are directly modeled and corrected for. We have begun to do this with AlphaPeptDeep (Zeng *et al*, 2022), which we also used to derive the predicted DIA libraries.

While proteome depth is not compromised by dimethyl labeling per se, we did notice a drop in identifications for multiplexed as opposed to single proteomes (although overall identifications and quantifications are still vastly higher). As this correlates with the number of labeled channels, we attribute it to the increased complexity of the DIA data which will eventually lead to an overlap in DIA transitions. At least some of this drop in identifications should be retrievable by further optimization of the DIA software as the extra complexity follows a clearly defined pattern in the channel spacing. In tri-plex format and combined with the relatively short gradients on the Evosep instruments, this achieves a throughput of 3 x 60 samples per day for proteome measurements, which would make it quite practical to incorporate an up-front fractionation step in the future.

We also explored the idea of a reference channel in single-cell mDIA. Note that this is conceptually different from the booster channel employed in the SCoPE-MS method (Budnik *et al*, 2018) because the fragments of the mDIA channels are offset from each other and do not contribute to a common low reporter mass. Existing DIA software such as Spectronaut or DIA-NN, which we used here, do not explicitly make use of the reference channel yet, and the concept of a separate FDR for individual target channels is not fully developed yet. Nevertheless, we found that identifications roughly doubled, from just above thousand proteins for interphase cells in our previous single-cell publication (Brunner *et al*, 2022) to a median of 2,370 now. Encouragingly, almost 4,000 proteins were identifiable for some of the cells. When combined with further ongoing instrumental and sample preparation advances, this may enable characterization of more than half of the expressed proteome in single cells in the near future. For our single-cell analysis, we use the low flow, high sensitivity Whisper gradients on the Evosep, leading to a throughput of 80 single cells per day when using a reference channel, which would increase to 160 per day or 1000 per week with the Lys-N five-plex method (subtracting one channel for the reference).

We are currently exploring the applicability of mDIA for a wide variety of proteomics questions and projects. In regular proteomics analyses, it retains most of the advantages of DIA in general, while tripling the throughput at little if any extra cost, as others have found as well (Derks *et al*, 2022). In our Deep Visual Proteomics technology (DVP) (Mund *et al*, 2022), mDIA would be especially attractive because different cell states that are spatially separated can be directly compared to each other. Furthermore, we found that the exquisite sensitivity of mDIA with a reference channel now allows the analysis of true single cells even in a spatial context (Schober *et al*, 2022). We look forward to further exploration of the reference channel concept. Combined with the inherent simplicity of the DIA acquisition method, it may lead to much more complete proteomes. Furthermore, the reference channel will make proteomic results much more comparable across experiments and between laboratories, something that should greatly benefit the community.

## Material and Methods

### Reagents and Tools

Reagents and Tools table
| Chemicals, enzymes and other reagents |  |  |
| --- | --- | --- |
| Bovine Serum Albumin (BSA) | New England BioLabs Inc. | P8108S |
| HeLa tryptic digest | Pierce | 1862824 |
| Yeast (*S. cerevisiae*) tryptic digest | Promega | V746A |
| *E. coli* tryptic digest | Waters | 186003196 |
| Triethylammonium bicarbonate (TEAB) | Sigma Aldrich | T7408 |
| Ammonium bicarbonate (ABC) | Merck | 1.01131 |
| Urea | Sigma Aldrich | U1250 |
| Dithiothreitol (DTT) | Sigma Aldrich | D0632 |
| Chloroacetamide (CAA) | Sigma Aldrich | C0267 |
| Formaldehyde - 'light' (CH<sub>2</sub>O) | Sigma Aldrich | F8775 |
| Formaldehyde-d<sub>2</sub>-solution - 'intermediate' (CD<sub>2</sub>O) | Sigma Aldrich | 492620 |
| Formaldehyde-<sup>13</sup>C,d<sub>2</sub>-solution - 'heavy' (<sup>13</sup>CD<sub>2</sub>O) | Sigma Aldrich | 596388 |
| Sodium cyanoborohydride - 'light' (NaBH<sub>3</sub>CN) | Sigma Aldrich | 156159 |
| Sodium cyanoborodeuteride - 'heavy' (NaBD<sub>3</sub>CN) | Sigma Aldrich | 190020 |
| Lichropur ® ammonia solution, 25% (NH<sub>3</sub>) | Merck | 5.33003 |
| LC-MS-grade water | Fisher Chemical | W6-4 |
| Trifluoroacetic acid (TFA) | Sigma Aldrich | 8.08260 |
| Formic acid (FA) | Sigma Aldrich/Merck | #64-18-6 |
| Acetonitrile | Sigma Aldrich/Merck | #75-05-8 |
| CDS Empore C18 Extraction Disks | Empore | 13-110-018 |
| Trypsin | Sigma-Aldrich | T6567 |
| Lys-C | FUJIFILM Wako | 125-05061 |
| Lys-N | ImmunoPrecise Antibodies (Europe) BV | L101 |
| **Software** |  |  |
| MaxQuant (2.0.1) | https://maxquant.org/ | N/A |
| DIA-NN (1.8.1) | https://github.com/vdemichev/DiaNN | N/A |
| RStudio (2022.07.2) and R (4.2.2) | https://posit.co/ | N/A |
| Jupyter Notebook | https://jupyter.org/ | N/A |
| **Other** |  |  |
| 384-well plates | Eppendorf | #0030129547 |
| Adhesive PCR Sealing Foil Sheets | Thermo Scientific | #AB-0626 |
| Bravo robot | Agilent |  |
| ThermoMixer | Eppendorf | #460-0223 |
| Mastercycler X50h | Eppendorf | #63160000 |
| NanoDrop One/OneC Microvolume UV-Vis Spectrophotometer | Thermo Fisher | @ND-ONEC-W |
| EvoSep One | EvoSep | #EV-1000 |
| Evotip Pure | EvoSep | #EV-2011 |
| timsTOF HT | Bruker Daltonik GmbH | N/A |
| timsTOF SCP | Bruker Daltonik GmbH | N/A |
| Column oven | Sonation lab solutions | #PRSO-V2 |
| Aurora Elite CSI | IonOpticks | AUR3-15075C18-CSI |

### Sample preparation of bulk samples

#### Bovine Serum Albumin

CAM-modified tryptic digest of bovine serum albumin (New England BioLabs Inc.) was labeled with dimethyl using the in-solution labeling protocol (Boersema *et al*, 2009). Briefly, 500 pmol of the BSA digest was reconstituted in 300 µL of 100 mM triethylammonium bicarbonate (TEAB, pH 7) buffer and aliquoted into three separate tubes. To each of these, 4 µL of 4% (vol/vol) formaldehyde (CH_2_O, CD_2_O or ^13^CD_2_O) and 4 µL of 600 mM sodium cyanoborohydride (NaBH_3_CN or NaBD_3_CN) were added sequentially. The samples were incubated at room temperature for 1 hour on a bench-top mixer. To quench the reaction, 16 µL of 1% (vol/vol) ammonia (NH_4_OH) was added to each tube. The solutions were then dried at room temperature using a speed vacuum concentrator, reconstituted in 200 µL of buffer A (0.1% formic acid), and desalted using in-house prepared, C18 StageTips (Rappsilber *et al*, 2003). The solutions were then adjusted to have a concentration of 50 fmol/μL. Three different mixtures were prepared by combining the three channels (Δ0/Δ4/Δ8) in varying mixing ratios: Mix 1 (8.5 μL / 1.0 μL / 0.5 μL), Mix 2 (7 μL / 2 μL / 1 μL), and Mix 3 (5 μL / 3 μL / 2 μL). A 1 μL aliquot of each mix was injected into the Thermo Orbitrap Exploris 480™ in triplicates, corresponding to an injection amount of 50 fmol per replicate.

#### Tryptic HeLa

A similar in-solution labeling protocol was followed for tryptic HeLa digests (Pierce #1862824) using 10 μg of starting material per channel. After C18 desalting, four different mixes were prepared: 62.5 ng/μL of label-free solution, 62.5 ng/μL of 1-channel (Δ0), 125 ng/μL of 2-plex (1:1 Δ0/Δ4), and 187.5 ng/μL of 3-plex (1:1:1 Δ0/Δ4/Δ8). A 2 μL aliquot of each mix was injected into the Thermo Orbitrap Exploris 480™ in triplicates and another 2 μL aliquot of each mix was injected into the Bruker timsTOF HT in triplicates. These correspond to injection amounts of 125 ng for unlabeled and one-channel solutions, 250 ng for two-plex solutions, and 375 ng for three-plex solutions, per replicate.

#### Mixed Species (Human / Yeast / E.coli)

For the mixed species experiment, three different mixtures with varying mixing ratios of HeLa tryptic digest (Pierce #1862824), *S. cerevisiae* tryptic digest (Promega V746A), and *E. coli* tryptic digest (Waters #186003196) were prepared prior to labeling: Mix 1 (3.25 µg / 0.75 µg / 1.00 µg Human(H)/Yeast(Y)/E.coli(E)), mix 2 (3.25 µg / 1.50 µg/ 0.25 µg H/Y/E), and mix 3 (3.25 µg / 0.25 µg / 1.50 µg H/Y/E). Mix 1 was then labeled with Δ0, mix 2 with Δ4, and mix 3 with Δ8, following the in-solution labeling protocol. After C18 desalting, each mixture was adjusted to 100 ng/µL using buffer A (0.1% formic acid) and the three labeled mixtures were combined with each other in a 1:1:1 ratio. A 3 μL aliquot of each mixture was injected into the Bruker timsTOF HT in triplicates for each of the two acquisition methods (20S and 16S4MS1, S = dia-PASEF scan) tested in this study. This corresponds to an injection amount of 300 ng total per replicate for each acquisition method.

#### Lys-N-derived HeLa

The Lys-N digest was prepared in-house. A HeLa cell pellet was lysed in 8.0 M urea, 50 mM ABC pH 8.5 and incubated in a Bioruptor for 10 minutes. Reduction of disulfide bonds was performed by adding dithiothreitol (DTT) to a final concentration of 10 mM DTT and incubating at room temperature for 45 minutes. Carbamidomethylation of the free sulfhydryl groups was achieved by adding chloroacetamide (CAA) to a final concentration of 40 mM CAA and incubating in the dark at room temperature for 15 minutes. The reaction was then quenched with DTT to a final concentration of 10 mM DTT. The resulting lysate was diluted to ensure that the concentration of urea in the solution is less than 2.0 M (diluted to 1.0 M). Lys-N (ImmunoPrecise Antibodies) was added to the lysate in a 1:100 (enzyme/protein) ratio and the digestion was left to digest at 37°C overnight. The mixture was acidified to a final concentration of 1% TFA to quench the digestion and desalted using in-house prepared, C18 stage tips. The same in-solution labeling protocol was used for labeling 25 μg of HeLa Lys-N peptides per channel. Five different mixes were prepared from the labeled peptides: 50 ng/μL of unlabeled solution, 50 ng/μL of one-channel (Δ0), 100 ng/μL of two-plex (1:1 Δ0/Δ4), 150 ng/μL of three-plex (1:1:1 Δ0/Δ4/Δ8), and 250 ng/uL of five-plex (1:1:1:1:1 Δ0/Δ2/Δ4/Δ6/Δ8). A 2 μL aliquot of each mix was injected into the Bruker timsTOF HT in triplicates. These correspond to injection amounts of 100 ng for unlabeled and one-channel solutions, 200 ng for two-plex solutions, 300 ng for three-plex solutions, and 500 ng for 5-plex solutions, per replicate.

### Sample preparation for reference channel and single-cell equivalents

HeLa cells were cultured in Dulbecco’s modified Eagle’s medium at 10% fetal bovine serum, 20 mM glutamine and 1% penicillin-streptomycin. The supernatant was carefully removed, cells were detached by accutase treatment for 5 min at 37°C followed by pipetting for cell aggregate dissociation. Cells were washed three times with ice-cold PBS, lysed by 60 mM TEAB and 20% acetonitrile (ACN) at 72°C for 30 min. Proteins were digested by trypsin and Lys-C in a 1:50 enzyme:protein ratio. Peptides were cleaned up by C18 desalting as described above. Cleaned up peptides were labeled as described above in 100 µL 60 mM TEAB (pH 8.5) to be comparable to the single-cell workflow. Evotips were loaded as described in the single-cell experiments.

### Sample preparation for single-cell experiments

HeLa cells were cultured and harvested following standard protocol as above. Cells were washed three times with ice-cold PBS, pelleted by centrifugation and resuspended in ice-cold PBS to a solution of 2x106 cells per mL. 2 µL propidium iodide (BioRad, 1351101) was added to the single-cell suspension and sorting was performed on the PI-negative live cell population using fluorescent-activated cell sorting (FACS). Single cells were sorted into 384-well plates containing 1 µL of 20% ACN, 0.01% n-Dodecyl β-D-maltopyranoside (DDM), 60 mM TEAB pH 8.5, centrifuged briefly, sealed with aluminum seal sheets and frozen at -80 °C until further processing. Single cells were incubated in the 384-well plate for 30 min at 72 °C in a PCR cycler with a lid temperature of 110 °C, digested overnight at 37 °C (lid temperature 55 °C) after addition of 1 µL of 20% ACN, 0.01% DDM, 60 mM TEAB and 0.5 ng trypsin and Lys-C each. Digested peptides were derivatized with dimethyl by adding 1 µL 0.6% formaldehyde (final concentration 0.15%) and 1 µL 92 mM cyanoborohydride (final concentration 23 mM). Single cells in a consecutive row pattern (B, D, F, …) were labeled with intermediate dimethyl (Δ4) using intermediate formaldehyde (CD_2_O) and light cyanoborohydride, while other wells (C, E, G, …) were labeled with heavy dimethyl (Δ8) using heavy formaldehyde (^13^CD_2_O) and heavy cyanoborohydride. After one hour incubation at room temperature, the dimethyl labeling reaction was stopped by adding 1 µL 0.65% ammonia solution (final concentration 0.13%). Before Evotip loading, samples were acidified by 1 µL 6% TFA (final concentration 1%). All pipetting steps were done with a Bravo robot (Agilent). Evotips Pure were loaded with the Bravo robot (Agilent), by activation with 1-propanol, washing two times with 50 µL Buffer B (99.9% ACN, 0.1% FA), activation with 1-propanol and two wash steps with 50 µL buffer A (99.9% H2O, 0.1% FA). In between Evotips were spun at 700xg for 1 min. For sample loading, Evotips were prepared with 70 µL buffer A and a short spin at 700xg. 10 µL of 1 ng/µL prepared and aliquoted reference channel proteome (labeled with dimethyl light, Δ0) was pipetted into Evotips, followed by an intermediate and heavy dimethyl labeled single cell with the same tip of the Bravo robot to avoid plastic contacts. Evotips were washed ones with 50 µL buffer A and stored at 4°C with buffer A on top until measured.

### Acquisition of bulk data

All samples were injected using the Thermo easyLC 1200 and separated via in-house pulled columns (50 cm length, 75 μm ID) packed with 1.9 μm ID ReproSil-Pur 120 C18-AQ (Dr. Maisch GmbH). These samples were sprayed into either an Orbitrap Exploris 480™ (Thermo Scientific) or a timsTOF HT (Bruker Daltonics) for MS analysis. Buffer A is 0.1% formic acid in LC–MS-grade water. Buffer B is 80% acetonitrile in LC–MS-grade water with 0.1% formic acid.

#### Bovine Serum Albumin

For BSA samples, a 30-min active gradient was used as follows: 2-7% Buffer B (minutes 0-1), 7-30% Buffer B (minutes 1-15), 30-65% Buffer B (minutes 15-18), 65-95% Buffer B (minutes 18-21), 95% Buffer B (minutes 21-24), 95-5% Buffer B (minutes 24-27), and 5% Buffer B (minutes 27-30). The flow rate was kept constant at 300 nL/min.

#### DDA

For determining the labeling efficiency of each dimethyl channel, shotgun DDA was employed on the Thermo Orbitrap Exploris 480™. The MS1 full scan range was 300–1,650 m/z with a resolution of 60,000, normalized automatic gain control (AGC) target of 300%, and a maximum injection time of 25 ms. This shotgun DDA approach was filtered for an intensity threshold of 5,000, with inclusion of charge states 2-5 and a dynamic exclusion duration of 30 s. There are 5 data dependent MS2 scans with a resolution of 15,000, normalized AGC target of 100%, maximum injection time of 28 ms, 1.4-Th isolation window width, and 27% normalized HCD collision energy. *DIA.* The DIA mode consists of one MS1 full scan followed by 16 MS2 windows with variable widths. Each MS1 full scan (300–1,650 m/z) was conducted at 120,000 resolving power, 300% normalized AGC target, and 100 ms maximum injection time. Each MS2 scan was conducted at 30,000 resolving power, 3,000% normalized AGC target, and 30% normalized HCD collision energy with a default charge of 2. The RF lens was set to 40%.

#### Tryptic HeLa

For the tryptic HeLa digests, a 75-min active gradient was used as follows: 2-7% Buffer B (minutes 0-1), 7-30% Buffer B (minutes 1-60), 30-50% Buffer B (minutes 60-66), 50-60% Buffer B (minutes 66-70), 60-90% Buffer B (minutes 70-71), and 90% Buffer B (minutes 71-75). The flow rate was kept constant at 300 nL/min. *Orbitrap DDA.* For determining the labeling efficiency of each dimethyl channel, shotgun DDA was employed on the Thermo Orbitrap Exploris 480™. The MS1 full scan range was 300–1,650 m/z with a resolution of 60,000, normalized automatic gain control (AGC) target of 300%, and a maximum injection time of 25 ms. This shotgun DDA approach was filtered for an intensity threshold of 100,000, with inclusion of charge states 2-5 and a dynamic exclusion duration of 30 s. There are 15 data dependent MS2 scans with a resolution of 15000, normalized AGC target of 100%, maximum injection time of 28 ms, 1.4-Th isolation window length, and 30% normalized HCD collision energy. *Orbitrap DIA.* The DIA mode consists of one MS1 full scan followed by 44 MS2 windows with variable widths (Steger *et al*, 2021). Each MS1 full scan (300–1,650 m/z) was conducted at 120,000 resolving power, 300% normalized AGC target, and 100 ms maximum injection time. Each MS2 scan was conducted at 30,000 resolving power, 3,000% normalized AGC target, and normalized HCD collision energy in stepped mode of 25%, 27.5%, and 30%. The default charge was 3 and the RF lens was set to 40%. *timsTOF dia-PASEF.* For the dia-PASEF measurements on the Bruker timsTOF HT platform, one MS1 full scan was followed by 20 dia-PASEF scans with variable widths that were optimized for the precursor densities of tryptic HeLa digests using py_diAID (Skowronek *et al*, 2022). The method covers an m/z range from 300 to 1,200 with two IM windows per dia-PASEF scan ranging from 0.3 to 1.7 Vs cm^-2^ (Fig EV4A). Since the MS1 scan and each dia-PASEF scan measures 100 ms, the total cycle time for this method is 2.1 s. The collision energy is a linear ramp from 20 eV at 1/K_0_ = 0.6 Vs cm^-2^ to 59 eV at 1/K_0_ = 1.6 Vs cm^-2^.

#### Mixed Species Experiment (Human / Yeast / E. coli)

The mixed species experiment was performed using the same gradient as the one used for tryptic HeLa digests. They were mainly measured in dia-PASEF mode on the Bruker timsTOF HT platform using two methods, hereby referred to as MS2-centric method (20S) and MS1-centric method (16S4MS1). The MS2-centric method has the same settings as the one described for tryptic HeLa digests where one MS1 scan is followed by 20 dia-PASEF scans (Fig EV4A). The MS1-centric method was also optimized using py_diAID to cover an m/z range from 300 to 1200 with two IM windows per dia-PASEF scan ranging from 0.3 to 1.7 Vs cm^-2^ (Fig EV4B). In this method, there are 4 ⨉ (1 MS1 scan + 4 dia-PASEF scans) that also leads to a total cycle time of 2.0 s (Fig EV4B). The collision energy is a linear ramp from 20 eV at 1/K_0_ = 0.6 Vs cm^-2^ to 59 eV at 1/K_0_ = 1.6 Vs cm^-2^.

#### Lys-N-derived HeLa

All Lys-N-derived HeLa measurements were acquired on the Bruker timsTOF HT using the 75-min gradient used for the tryptic HeLa digests. They were mainly measured in dia-PASEF mode using an MS2-centric method (20S) optimized for Lys-N-derived HeLa peptides (Fig EV6A).

### LC-MS/MS analysis of ultra-high sensitivity and single-cell data

All samples were loaded onto Evotips Pure and measured with the Evosep One LC system (EvoSep) coupled to a timsTOF SCP mass spectrometer (Bruker). The Whisper40 SPD (samples per day) method was used with the Aurora Elite CSI third generation column with 15 cm and 75 µm ID (AUR3-15075C18-CSI, IonOpticks) at 50 °C inside a nanoelectrospray ion source (Captive spray source, Bruker). The mobile phases comprised 0.1% FA in LC–MS-grade water as buffer A and 99.9% ACN/0.1% FA as buffer B. The timsTOF SCP was operated in dia-PASEF mode with variable window widths. Optimal dia-PASEF methods cover the precursor cloud highly efficient in the m/z – ion mobility (IM) plane while providing deep proteome coverage. For method generation with py_diAID, the precursor density distribution in m/z and IM was estimated based on a tryptic 48 high-pH fraction library (Skowronek *et al*, 2022). We calculated the optimal cycle time based on the chromatographic peak width of 5 ng HeLa single runs. DIA-NN reported a base-to-base peak width of 7.6 s, translating into 8 data points per peak at a cycle time of 0.96 s. The optimal dia-PASEF method consisted of one MS1 scan followed by eight dia-PASEF scans with two IM ramps per dia-PASEF scan, covering a m/z range from 300 to 1200 and IM of 0.7 to 1.3 Vs cm^-2^ (Fig EV6B). The mass spectrometer was operated in high sensitivity mode, the accumulation and ramp time was specified as 100ms, capillary voltage was set to 1400 V and the collision energy was a linear ramp from 20 eV at 1/K_0_ = 0.6 Vs cm^-2^ to 59 eV at 1/K_0_ = 1.6 Vs cm^-2^.

The label efficiency of 1 ng HeLa peptides was assessed on the timsTOF SCP operating in dda-PASEF mode with ten PASEF/MSMS scans per topN acquisition cycle. Singly charged precursors were excluded by their position in the m/z-IM plane using a polygon shape, and precursor signals over an intensity threshold of 1,000 arbitrary units were picked for fragmentation. Precursors were isolated with a 2 Th window below m/z 700 and 3 Th above, as well as actively excluded for 0.4 min when reaching a target intensity of 20,000 arbitrary units. All spectra were acquired within a m/z range of 100 to 1,700. All other settings were as described above.

### Raw data analysis with MaxQuant for label efficiency assessment

MaxQuant version 2.0.1 was used to process triplicate to quintuplicate DDA data for calculating the labeling efficiency of each dimethyl channel. All dimethyl groups (Var DimethNter0, Var DimethLys0, Var DimethNter2, Var DimethLys2, Var DimethNter4, Var DimethLys4, Var DimethNter6, Var DimethLys6, Var DimethNter8, and Var DimethLys8) were first configured as variable modifications in MaxQuant. They were selected as variable modifications together with Oxidation (M) and Acetyl (Protein-N-term). For example, BSA samples labeled with dimethyl Δ0 were searched with Var DimethNter0, Var DimethLys0, Oxidation (M), and Acetyl (Protein-N-term) as variable modifications. Carbamidomethyl (C) was selected as a fixed modification for bulk but unselected for ultra-high sensitivity acquisitions, since peptides were not reduced and alkylated. The maximum number of modifications per peptide was set to 3. Depending on the sample, either trypsin/P or Lys-N was selected as the protease and the maximum number of missed cleavages was set to 2. Labeling efficiency was calculated based on intensity ratios of labeled peptides relative to all detected peptides.

### Raw data analysis with DIA-NN

DIA-NN version 1.8.1 was used to search DIA raw files and dia-PASEF files for precursor and fragment identifications based on 2D or 3D peak position (retention time, m/z precursor, and IM) using AlphaPeptDeep-predicted spectral libraries (*dimethyl labeled tryptic HeLa: 6331933 precursor entries, label-free tryptic HeLa: 6341464 precursor entries, dimethyl labeled tryptic Human/Yeast/E.Coli mix: 9716057 precursor entries, dimethyl labeled Lys-N-derived HeLa: 3387578 precursor entries, dimethyl labeled tryptic HeLa for ultra-high sensitivity samples: 6294089 precursor entries*).

The DIA-NN search included the following settings: Protein inference = “Genes”, Neural network classifier = “Single-pass mode”, Quantification strategy = “Robust LC (high precision)”, Cross-run normalisation = “RT-dependent”, Library Generation = “IDs, RT and IM Profiling”, and Speed and RAM usage = “Optimal results”. Mass accuracy and MS1 accuracy were set to 0 for automatic inference from the first run. The following settings were also enabled: “Use isotopologues”, “MBR”, “Heuristic protein inference”, and “No shared spectra”. For dimethyl labeled BSA and tryptic HeLa samples, the following additional commands were entered into the DIA-NN command line GUI: *(1)* {--fixed-mod Dimethyl, 28.0313, nK}, *(2)* {--channels Dimethyl, 0, nK, 0:0; Dimethyl, 4, nK, 4.0251:4.0251; Dimethyl, 8, nK, 8.0444:8.0444}, *(3)* {--original-mods}, *(4)* {--peak-translation}, *(5)* {--ms1-isotope-quant}, *(6)* {--report-lib-info}, and *(7)* {–mass-acc-quant 10.0}. Note that *(7)* is removed from the command line for the MS1-centric method. For dimethyl labeled Lys-N-derived HeLa samples, additional channels were inserted into the DIA-NN command line GUI: {--channels Dimethyl, 0, nK, 0:0; Dimethyl, 2, nK, 2.0126:2.0126; Dimethyl, 4, nK, 4.0251:4.0251; Dimethyl, 6, nK, 6.0377:6.0377; Dimethyl, 8, nK, 8.0444:8.0444}. For label-free HeLa tryptic/Lys-N samples, only the following commands were used: *(1)* {--original-mods}, *(2)* {--ms1-isotope-quant}, *(3)* {--report-lib-info} and *(4)* {–mass-acc-quant 10.0}. In cases where only one channel or two channels were present, the actual number of channels were inserted into the DIA-NN command line GUI and not the full set of channels We used reannotate with the fasta SwissProt database of reviewed sequences (April, 2022).

Ultra-high sensitivity measurements were searched in the same way except that the scan window was always set to a fixed value of 9, as well as MS1 and mass accuracy to 15 ppm, as this was identified as the optimal settings after several runs and kept constant for minimizing variance between the datasets. In cases of empty channels, the full set of channels were still added to the DIA-NN command line GUI. The single-cell dataset was searched with same settings against a subset of files (50 raw files) and then the generated report-lib library (file name: ‘report-lib.tsv’, from ‘first-pass’ search) was used to analyze all raw files. The report-lib library was adjusted by renaming the ‘Protein.Group’ column to ‘UniprotID’ and searched without the ‘reannotate’ function in DIA-NN to keep the same protein grouping. The first-pass-result of this search was used for data analysis, since a MBR cycle was already performed in the report-lib library generation.

### Protein quantification for bulk data

The DIANN R package was used to calculate the MaxLFQ abundance for protein groups (Demichev *et al*, 2019). The MaxLFQ abundance (Cox *et al*, 2014) was calculated based on the ‘Precursor.Translated’ column output by DIA-NN for MS2-centric methods whereas it is based on the ‘Ms1.Area’ column output by DIA-NN for the MS1-centric method. For all datasets, the output results from the R package were then filtered for ‘Global.PG.Q.Value’ < 0.01 and ‘PG.Q.Value’ < 0.05. An additional filtering for ‘Channel.Q.Value’ < 0.01 was applied for labeled datasets and ‘Q.Value’ < 0.01 for label-free datasets. For the mixed species three-plex sample, the calculated protein ratios were normalized using the human protein ratios which are present in equal amounts in the different channels.

### RefQuant and single-cell protein quantification

The filtered DIA-NN output report table was processed with the ‘RefQuant’ package implemented in Python. RefQuant imports the DIA-NN table and extracts the relevant information used for quantification (quantities used: ‘Fragment.Quant.Raw’, ‘Ms1.Area’, ‘Precursor.Translated’). Subsequently, on each precursor, the following operations are performed: The target quantity is divided by the reference quantity for each of the available extracted quantities. This results in a list of ratios. The ratios are sorted in ascending order and the first 40% of ratios are retained. The overall ratio R between target and reference is then obtained by taking the mean of the remaining ratios. The ratio R is then multiplied by a precursor-specific scaling factor, representing the intensity in the reference channel, which results in an overall intensity estimate of the precursor in the target channel. The scaling factor for each precursor is derived by taking the median of the reference intensity over all available runs for this particular precursor. The default value for the reference intensity is derived from the ‘Ms1.Area’. In case there are remaining precursors with less than two not-null ‘Ms1.Area’ values, the scaling factor is derived from ‘Precursor.Translated’. Otherwise, if there are still remaining precursors with less than two not-null reference intensity values, the scaling factor is derived from the summed available ion intensities. Processing with RefQuant resulted in a quantification matrix (protein group, precursors, experiment and channel RefQuant intensity).

The RefQuant quantification matrix was then further processed with the iq package (Pham *et al*, 2020) in order to derive MaxLFQ protein quantities.

### Proteomics downstream data analysis

Proteomics data analysis was performed in RStudio 2022.07.2 with R 4.2.2 and Python (version 3.8.2). MaxQuant output tables were filtered for ‘Reverse’, ‘Only identified by site modification’ and ‘Potential contaminants’ before further processing. For ultra-high sensitivity measurements, the RefQuant output was filtered for ‘Lib.PG.Q.Value’ < 0.01, ‘Q.value’ < 0.01 and ‘Channel.Q.Value’ < 0.15. 764 cells were measured and 534 cells were not empty (above 1% of identified proteins). Cells in a range of 1.5 standard deviations around the median number of identified proteins were used for further analysis, except three outliers which showed a highly increased MS signal intensity compared to their proteomic depth (SM03p5b_30_5329_label4, SM03p4a_68_5210_label8, SM03p6a_65_5473_label8).

### Single-cell protein and RNA comparison and dropout statistics

The SMART-Seq2 (Hu et al, 2019) dataset contained 720 HeLa cells in three different batches with a total of 24,990 expressed genes. The Drop-seq (Schwabe et al, 2020) dataset measured three batches with a total of 5,665 cells and 41,161 expressed genes. Single-cell analysis was performed with scanpy v1.6.0 (Wolf et al, 2018) similar as before, while proteins were normalized based on the sceptre ‘normalize’ method that iteratively adjusts the median protein expression between the different batches and different targets (Schoof *et al*, 2021). If not stated otherwise, standardized filtering across all datasets, removed cells with less than 600 genes expressed, and removed genes detected in < 15% of the remaining cells, resulting in 10,557 transcripts in 720 cells in the SMART-Seq2 dataset and 5,022 transcripts and 6,701 cells measured with Drop-seq technology. The data completeness across covered dynamic range was computed as a function of the mean log(x + 1)-transformed protein abundance of all non-zero/-NaN entries. We included the expected data completeness based on the assumption that missing values are purely due to shot-(Poisson)-noise as 1-exp(-x). For correlation analysis, the RNA abundance entries were linearly scaled to sum to the mean cell size of the respective dataset per cell (231,281.56 for SMART-Seq2 and 7,808.12 for Drop-Seq) followed by log(x + 1) transformation of all abundance entries. Entries of missing protein abundance values were excluded from the specific computation. In coefficient of variation (CV) versus mean intensity plots comparing different technologies as well as the mean versus CV analysis (including the core proteome analysis) and the CV distribution boxplots, RNA expression vectors were scaled to the mean cell size of that measurement technology. Mean and CV values were computed per gene under the assumption that single-cell RNA-sequencing data are not zero inflated (Svensson, 2020) while NaNs were excluded for the proteomics data. CV (Proteomics) versus CV (RNA-seq) plots show the comparison of CV values of proteins/genes that were shared between all datasets.

## Supplemental data

This article contains supplemental data.

## Acknowledgement

This study was supported by the Max Planck Society for Advancement of Science, European Union’s Horizon 2020 research and innovation program under grant agreement No 874839 (ISLET), the Deutsche Forschungsgemeinschaft project “Chemical proteomics inside us” (grant no.: 412136960) and by the Bavarian State Ministry of Health and Care through the research project DigiMed Bayern (www.digimed-bayern.de). MT and MW acknowledge support from the International Max Planck Research School for Life Sciences – IMPRS-LS. SR is supported by the Helmholtz Association under the joint research school "Munich School for Data Science” -MUDS. FAS is an EMBO postdoctoral fellow (ALTF 399-2021). We are grateful for the FACS support by the Imaging Core Facility at the Max Planck Institute of Biochemistry, in particular Martin Spitaler and Markus Oster. We thank our colleagues in the Department of Proteomics and Signal Transduction, Max Planck Institute of Biochemistry and at the Center for Protein Research at Copenhagen University, for discussions and support. In particular, we thank I. Paron and T. Heymann for technical support and column production and Medini Steger for scientific administration support.

## Author contributions

MT, MS and MM conceptualized and designed the study. MT, CEMI, FAS, MW, PS and MS performed experiments. CA developed quantification software. SR and FJT conceptualized the single-cell modeling. W-FZ and X-XZ predicted AlphaPeptDeep libraries. MT, CEMI, CA, IB, FAS, MW, SR, A-DB, MS and MM analyzed the data. MT, CEMI and PS curated data. MT, MS and MM supervised the project. MT, CEMI, MS and MM wrote the original manuscript draft. All authors read, revised and approved the manuscript.

## Supplementary Figures

**Fig EV1:**
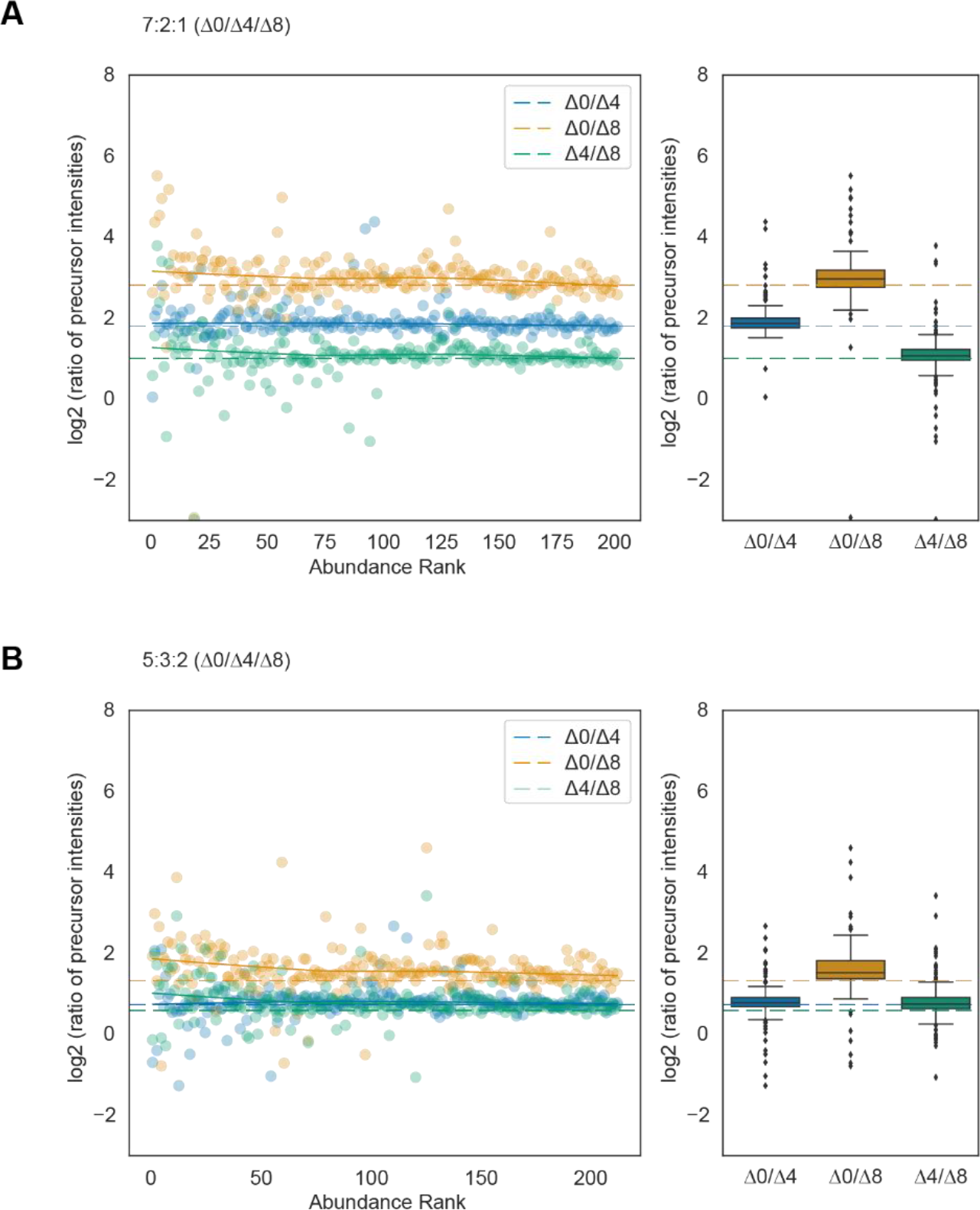
Quantification accuracy of bovine serum albumin (BSA)-derived tryptic peptides labeled with dimethyl mass tags and mixed at defined ratios. A and B. Quantification accuracy for peptides labeled with three mass tags and mixed in 7:2:1 (A) or 5:3:2 (B) ratios. Scatter plots on the left panel show the log2 intensity ratios as a function of the peptide abundance rank. In both panels, the expected ratios are marked by colored dashed lines.

**Fig EV2:**
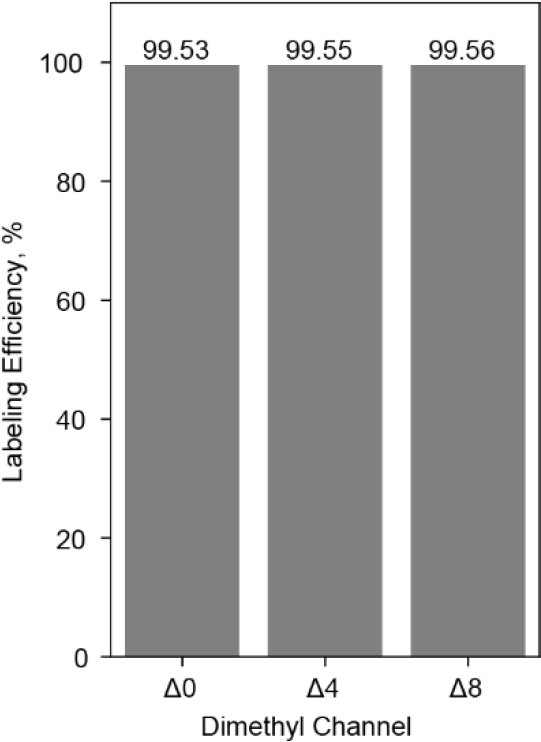
Labeling efficiency of tryptic HeLa peptides acquired on the Orbitrap platform. Tryptic peptides from HeLa cells were labeled with dimethyl mass tags Δ0, Δ4 and Δ8 and acquired individually in DDA mode. Labeling efficiencies based on intensity ratios of labeled peptides relative to all detected peptides are shown.

**Fig EV3:**
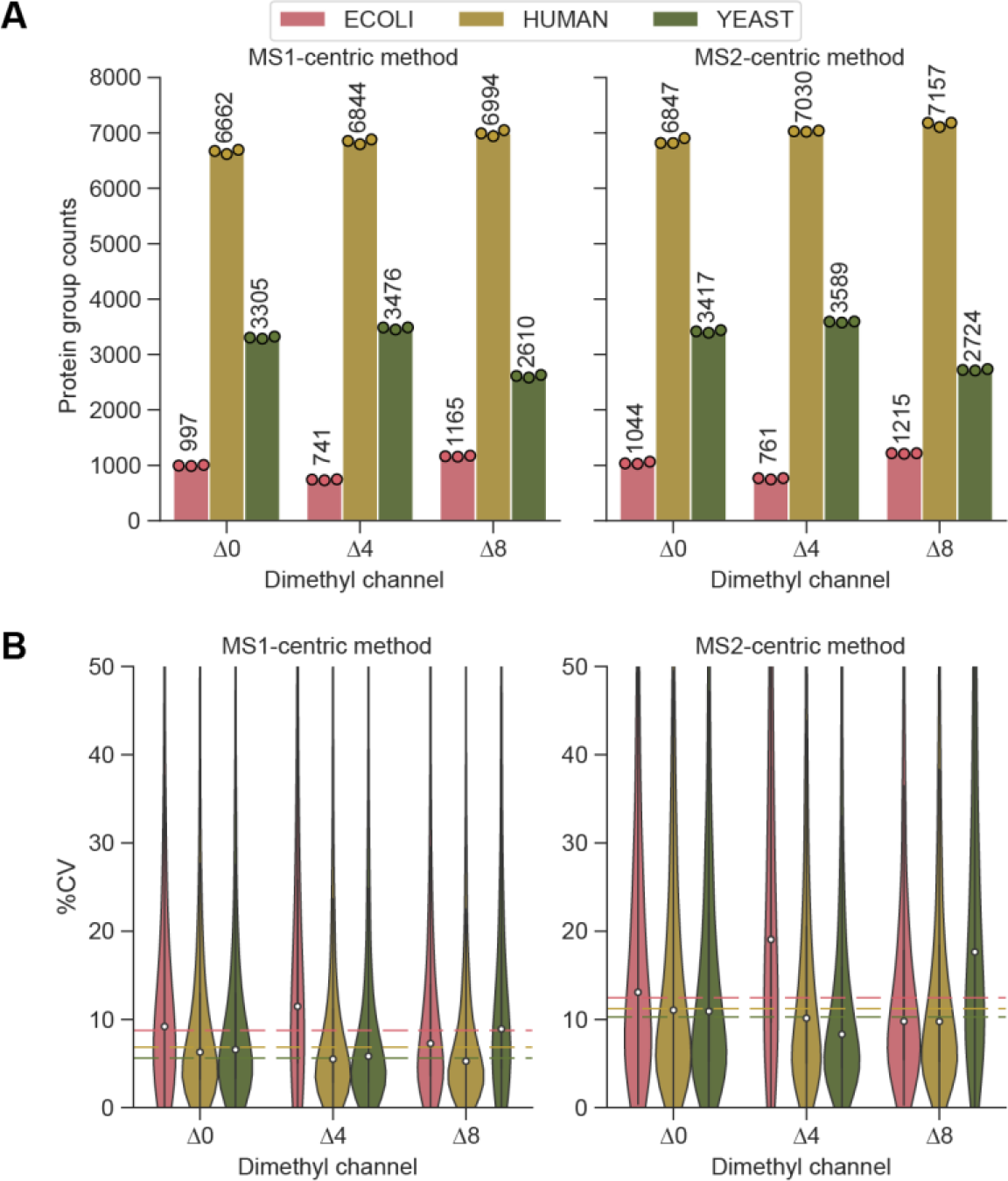
Identification rates and coefficient of variation (CVs) in the mixed species experiment. A. Tryptic peptides from E. coli, yeast and human were combined at defined ratios before dimethyl labeling. Side-by-side comparison of the number of quantified protein groups for the individual species with MS1- and the MS2-centric methods. 100 ng of peptides were injected per channel. B. Coefficients of variation (CVs, %) of protein groups shown in (A). Median CVs are shown as dashed lines.

**Fig EV4:**
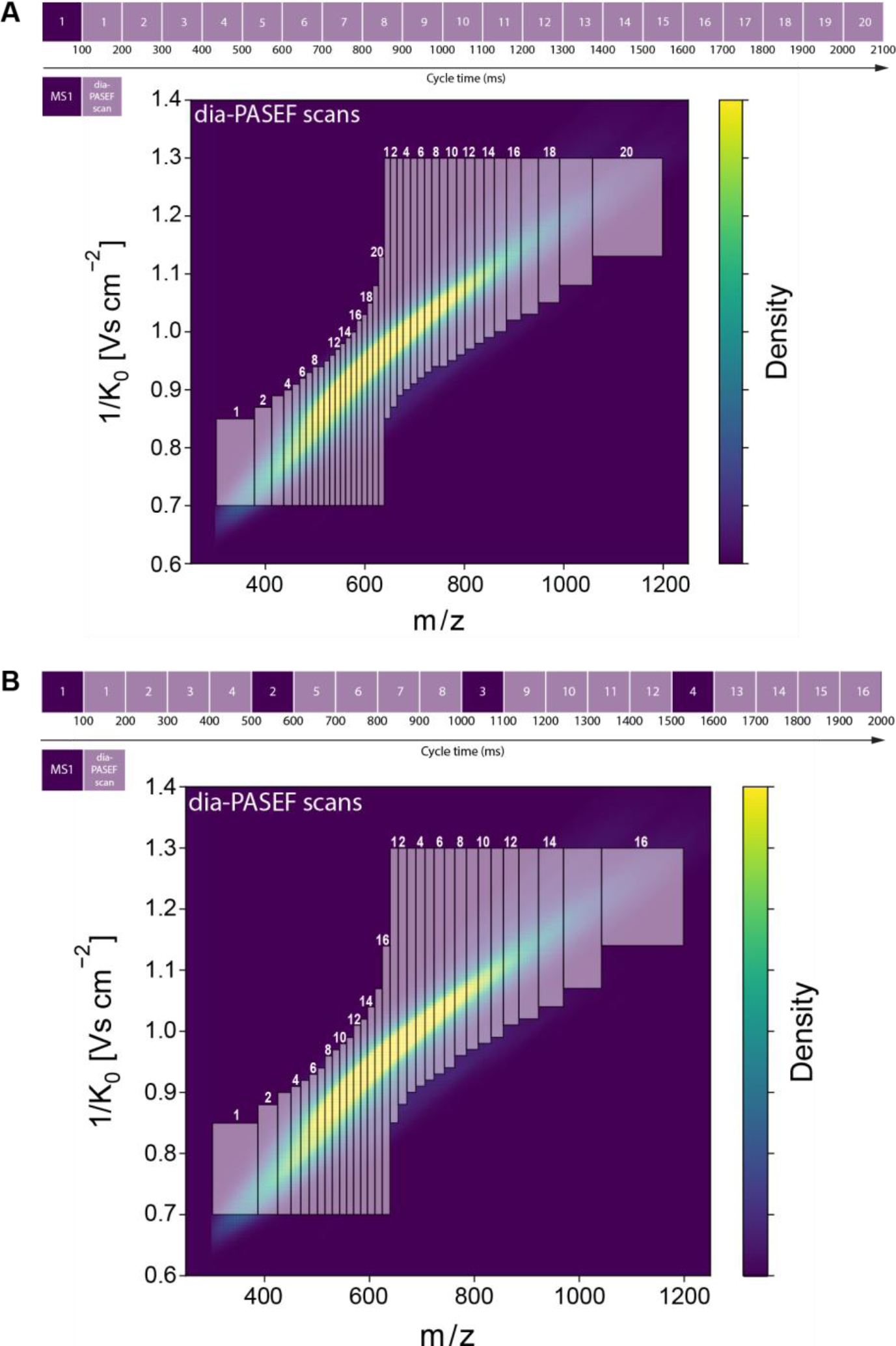
dia-PASEF acquisition methods optimized for dimethylated tryptic HeLa peptides. A. 20 dia-PASEF scan method used for MS2-centric acquisitions. It consists of one MS1 scan followed by 20 dia-PASEF scans with variable m/z isolation widths and two ion mobility windows in one cycle (∼2.1 seconds). B. 16 dia-PASEF scan method used for MS1-centric acquisitions. This method consists of 16 dia-PASEF and four MS1 scans, with each MS1 scan followed by 4 dia-PASEF scans. Similar to the 20 dia-PASEF scan method (A), the cycle time is about two seconds.

**Fig EV5:**
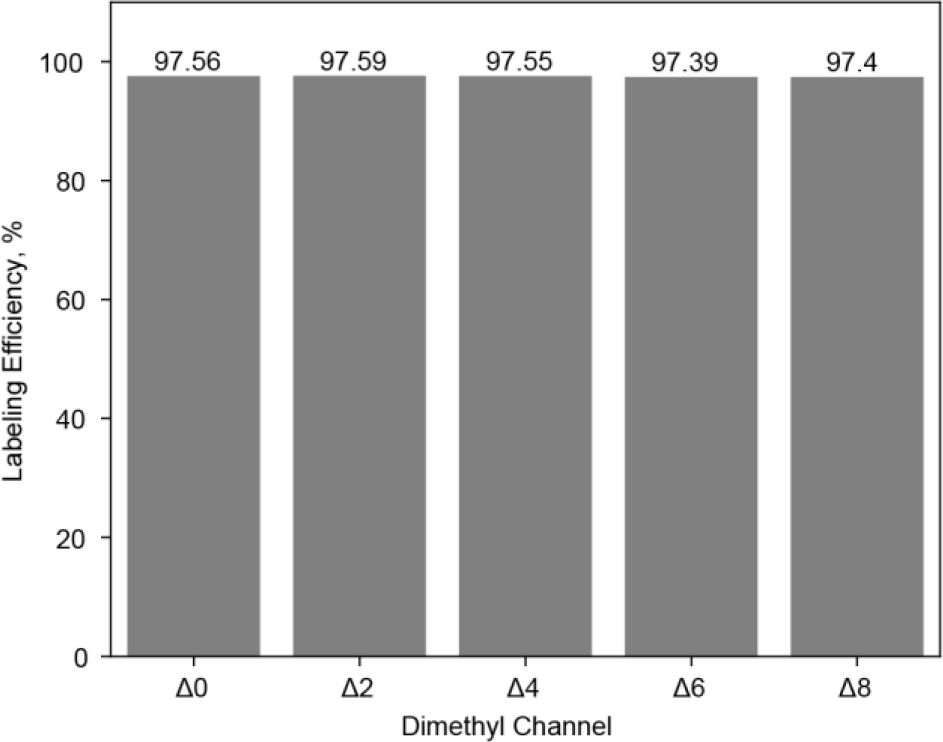
Labeling efficiency of Lys-N-derived HeLa peptides acquired on the Orbitrap platform. Lys-N-derived HeLa peptides from HeLa cells were labeled with dimethyl mass tags Δ0, Δ2, Δ4, Δ6, and Δ8 and acquired individually in DDA mode. Labeling efficiencies based on intensity ratios of labeled peptides relative to all detected peptides are plotted.

**Fig EV6:**
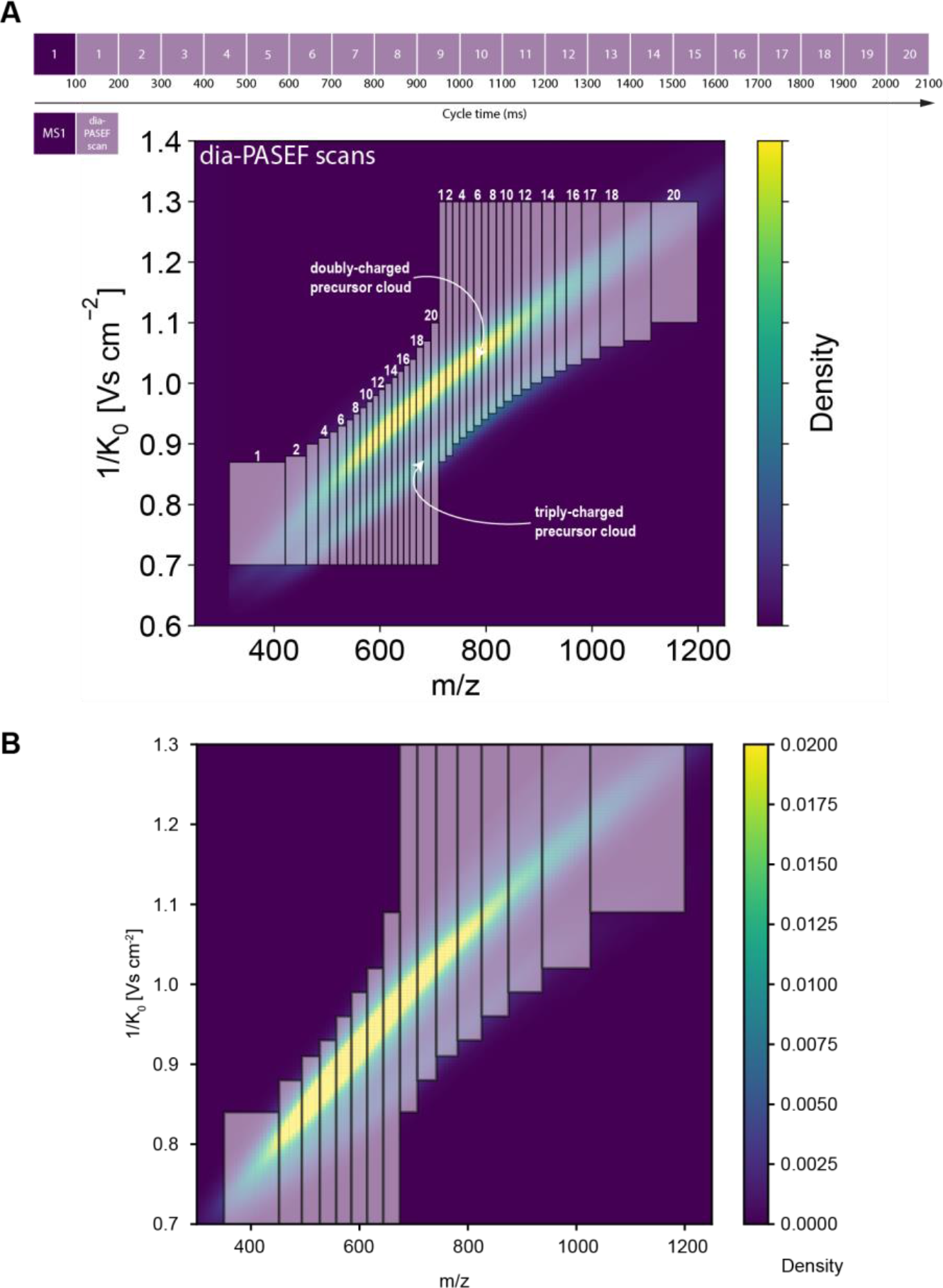
Optimal dia-PASEF acquisition method for Lys-N-derived HeLa peptides and tryptic HeLa for ultra-high sensitivity measurements. A. 20 dia-PASEF scan method was used for the acquisition of Lys-N-derived HeLa peptides. The method was optimized to specifically cover both doubly- and triply-charged precursors. One cycle time is about 2.1 seconds. B. Eight dia-PASEF scan method for ultra-high sensitivity measurement of single-cell equivalents and single cells using the ‘Whisper40 SPD’ gradient in combination with an Aurora Elite LC column (IonOpticks). The acquisition scheme is plotted on top of a kernel density distribution of the precursors from a representative tryptic 48 high-pH fraction HeLa library (Skowronek *et al*, 2022).

**Figure EV7:**
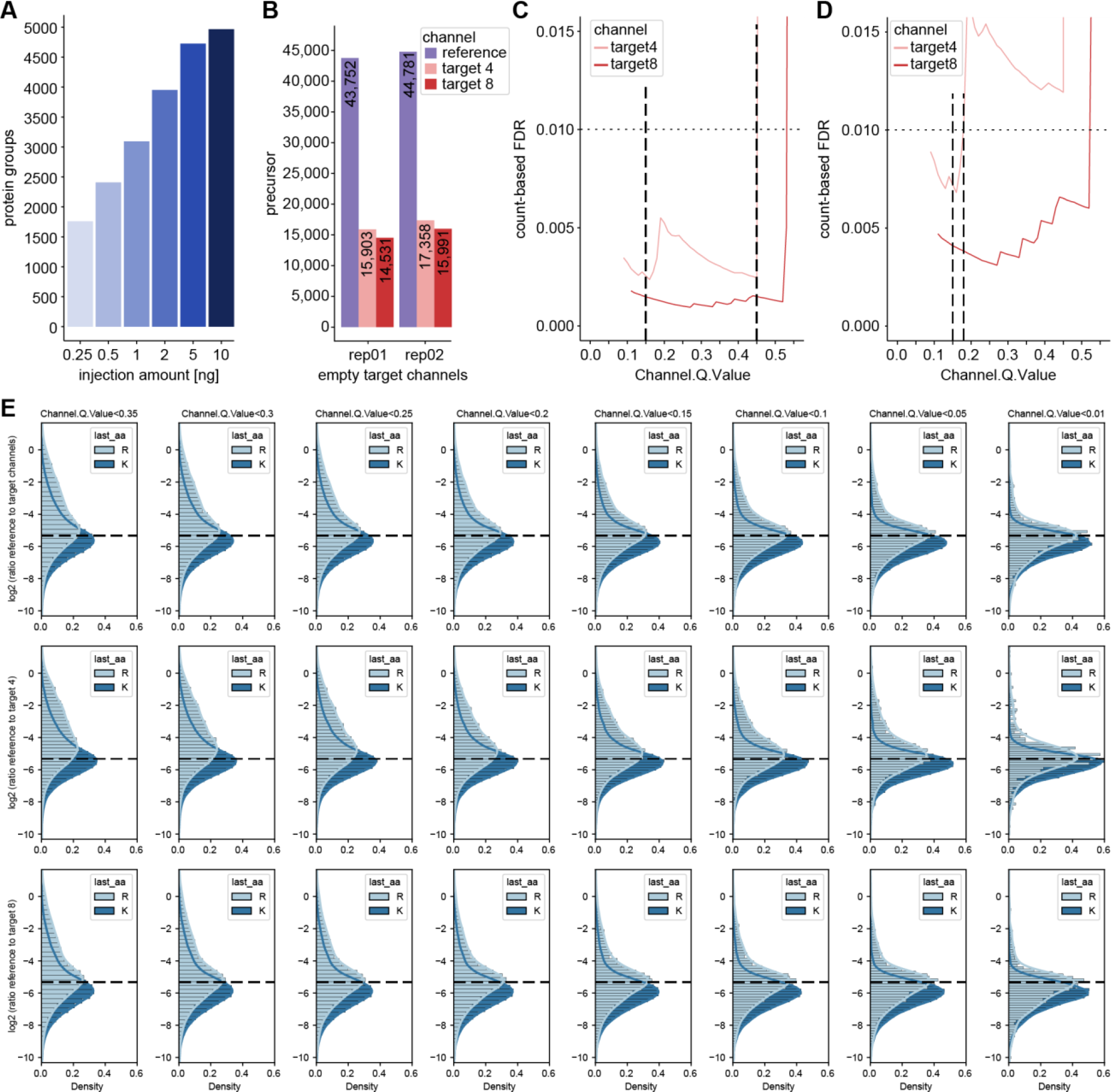
Evaluation of ‘Translated.Q.Value’ and ‘Channel.Q.Value’ on identifications in empty target channels and quantification in single-cell equivalents. A. Dilution series of HeLa peptides to define protein identifications in mDIA workflow, in which the linear increase with input amount levels off at about 10 ng. B. Precursor identifications of empty target channels at 1% ‘Translated.Q.Value’ revealed more than 1% false positives. C and D. Count-based FDR estimation on precursor (C) and protein level (D) by dividing two quantities, namely the maximum number of identified precursors or proteins in four empty target channel runs by the minimum number of identified precursors or proteins in a total of four target channels (target channel 4 and 8) which contain single-cell equivalent amounts. The count-based FDR reveals a ‘Channel.Q.Value’ filter of 0.45 for precursors and 0.17 for proteins at 1%. E. Quantification evaluation of ‘Channel.Q.Value’ filter. Ratios of reference to target channel (top), channel 4 (middle) and 8 (bottom) by RefQuant (see Fig 5) are shown for arginine and lysine precursor at a given filter of ‘Channel.Q.Value’. At all filtering steps, the modes of the distributions are close to the expected ratio (dashed line) with a systematic skew towards lower ratios, which is more pronounced for the arginine precursors. The skews in the distributions are decreased with increased filtering stringency in a gradual manner. The mean quantification ratios are as expected, whereas they start to deviate at 0.2.

**Figure EV8:**
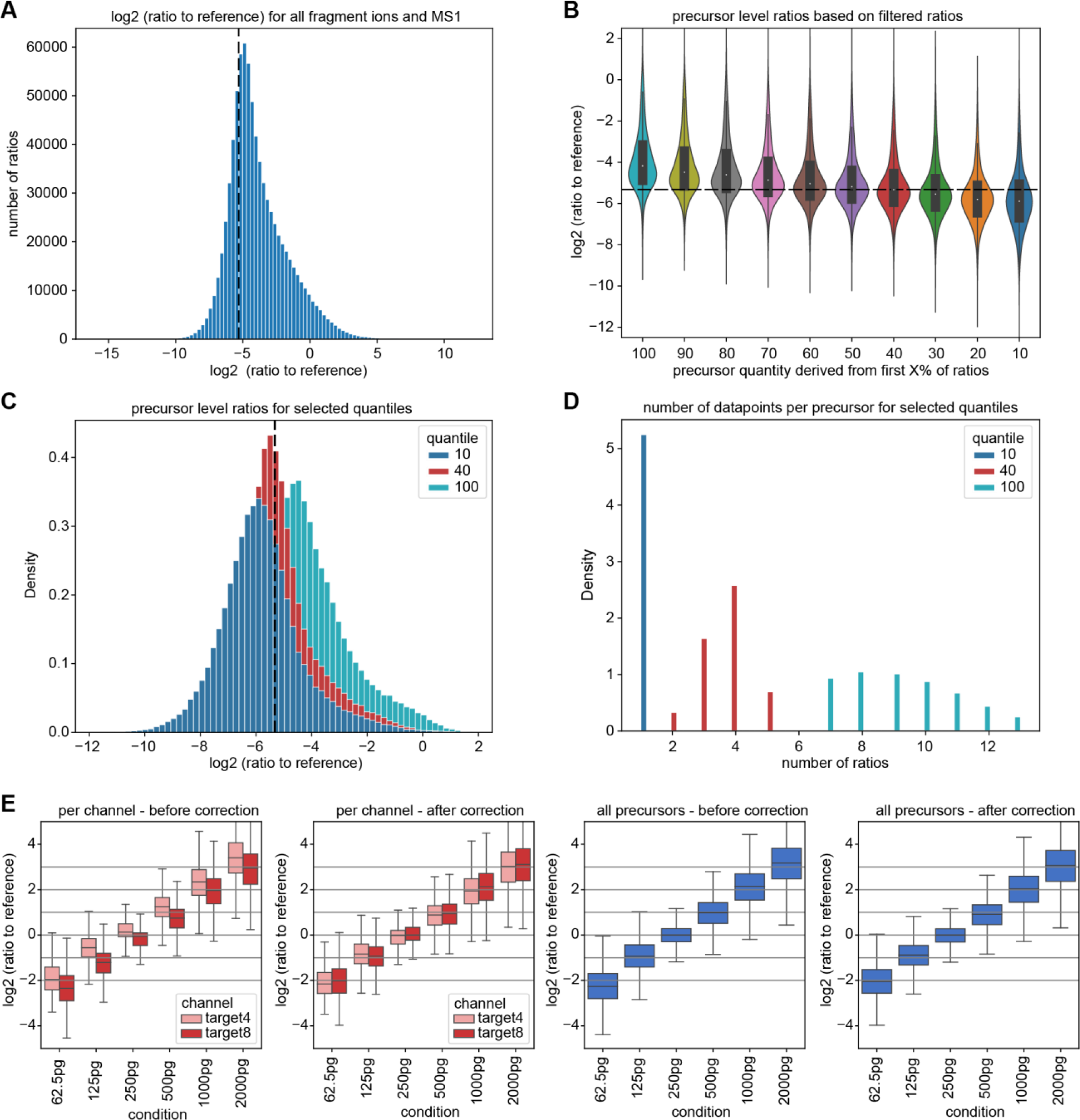
Ratios to reference channel evaluation and channel correction comparison in scQuant. A. Ratios to the reference for all available ions (fragment ions and MS1 peaks), revealing a distribution that has a mode proximate to the expected ground truth (dashed line). The distribution is asymmetric with a skew towards less extreme ratios. A potential explanation for this skew is that noise or interferences are dominant for a fraction of the ratios (a ratio of 0 might be a ‘noise vs. noise’ comparison). B. An approach to mitigate this asymmetry is to filter out some ions before estimating the precursor level ratio. For this, the ratios of a precursor are sorted ascending and only the first ratios are retained (up to the ‘X value’ indicated). The precursor ratio is estimated by taking the mean of the remaining ratios. We see that retaining 40% of the ratios matches the ground truth well and is more symmetric. C. A more detailed comparison of the distributions when taking 10% or 40% of ratios as compared to all ratios. D. Number of available ratios per precursor after filtering. We see that in general between 7 to 14 ratios are available, which reduces to 2 to 5 when taking the 40% quantile. E. Channel correction comparison in scQuant dataset in comparison of all precursor and target channel. Ratio of reference channel to target channel was calculated using RefQuant. All log2 ratios were normalized to the single-cell equivalent sample in the target channel (250 pg). A basic median normalization between the channels was applied.

## Supplemental Text EV9

Fundamental considerations about the noise in target and reference for ratio-based quantification

We investigated to what extent, if any, the ratio between reference and target channels would contribute to variance in the experimental values. If a given peptide had a 50% error in its estimated signal in a 40:1 reference to target ratio, this would manifest in a 20.5-fold ratio in a first run. In a second run – given that MS signals are very reproducible – we would then expect a ratio of 20 or perhaps 19.2. From this it follows that -concerning the noise - the variance of the ratio estimation over runs is not determined by magnitude of the ratio. Following this reasoning, the overall variance of the log2 ratios *σ*^2^_*target*_ is *σ*^2^_*ratio*_ = *σ*^2^_*ratio*_ + *σ*^2^_*refrence*_, with *σ*^2^_*ratio*_ denoting the variance of the target channel and *σ*^2^_*target*_ denoting the variance of the reference channel. As the higher abundances of the reference channel stabilizes its signal, we conclude that the overall variation of the ratios will decrease with higher reference proteome amounts.

**Figure EV10:**
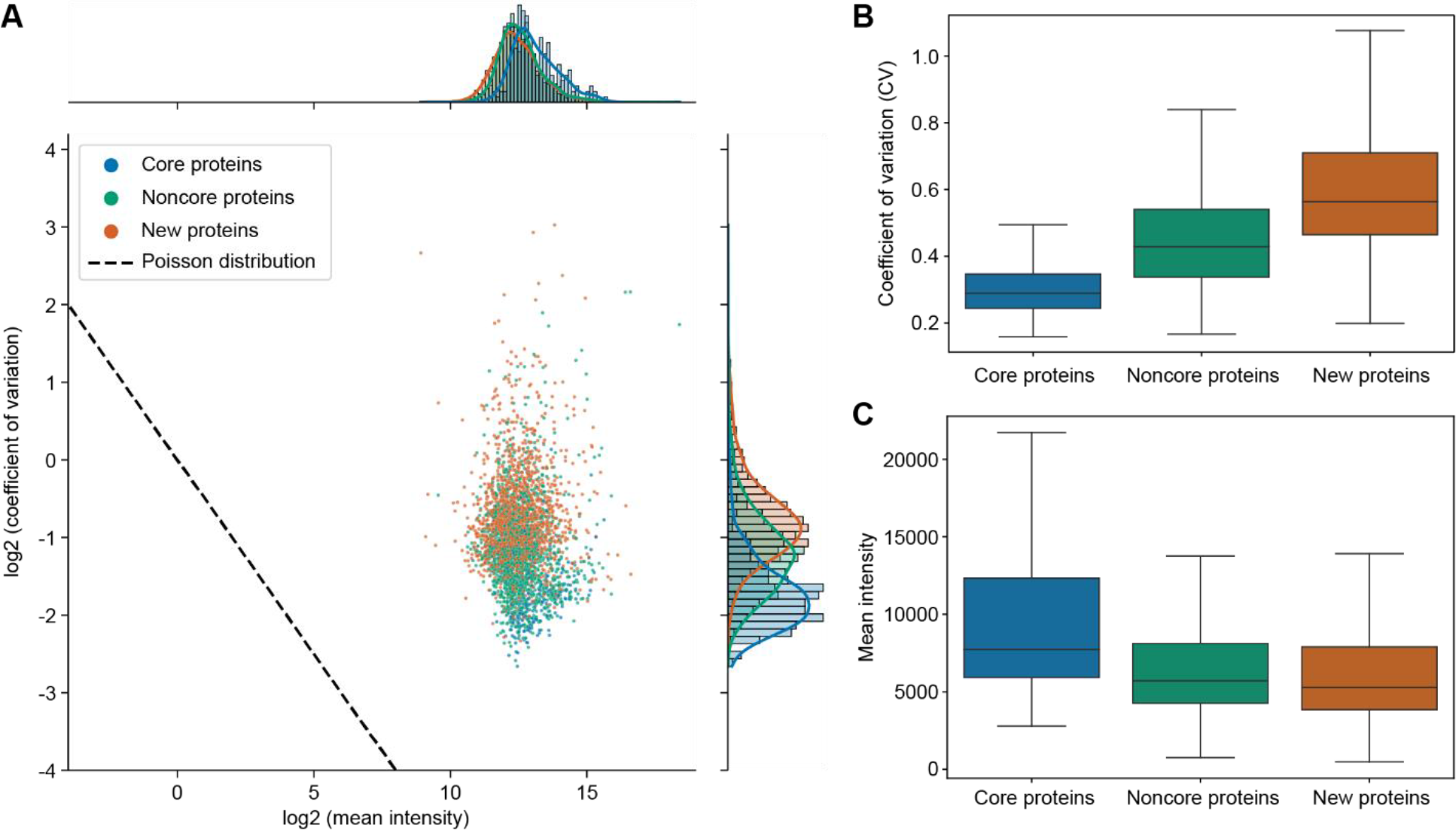
mDIA shows that the concept of a stable proteome is still valid at higher proteomic depth in single cells. A. Coefficients of variation of single-cell mDIA protein expression plotted against the mean intensity of each protein. ‘Core proteins’ are labeled in blue, ‘noncore proteins’ in green and additionally identified and quantified proteins compared to our previous single-cell publication in orange (Brunner *et al*, 2022). B. Distribution of coefficients of variation across the mDIA single-cell dataset for ‘core’ (blue), ‘noncore’ (green) and ‘new’ (orange) proteins. C. Distribution of the mean MS intensity across the mDIA single-cell dataset for ‘core’ (blue), ‘noncore’ (green) and ‘new’ (orange) proteins.

