## Supplementary Figures for "Robust dimethyl-based multiplex-DIA workflow doubles single-cell proteome depth via a reference channel"

**A**

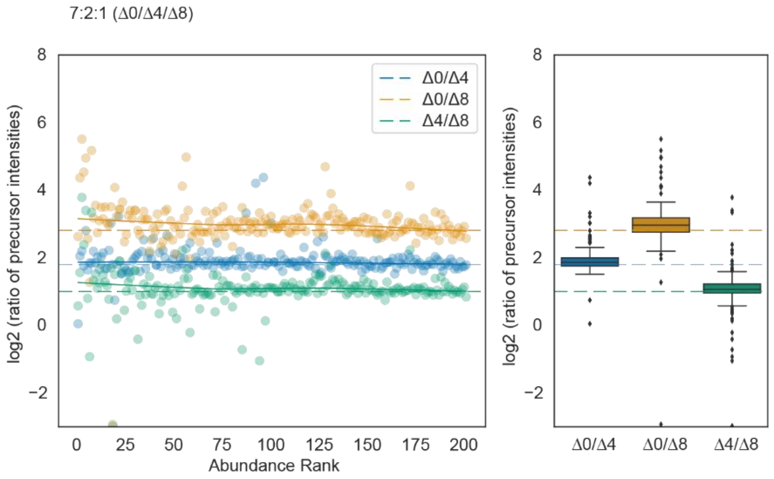


**B**

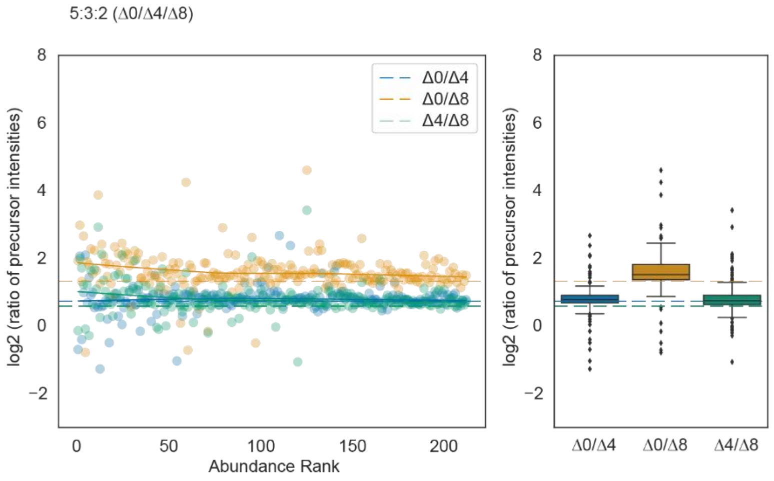


**Fig EV1: Quantification accuracy of bovine serum albumin (BSA)-derived tryptic peptides labeled with dimethyl mass tags and mixed at defined ratios.**

A and B. Quantification accuracy for peptides labeled with three mass tags and mixed in 7:2:1 (A) or 5:3:2 (B) ratios. Scatter plots on the left panel show the $log_2$ intensity ratios as a function of the peptide abundance rank. In both panels, the expected ratios are marked by colored dashed lines.

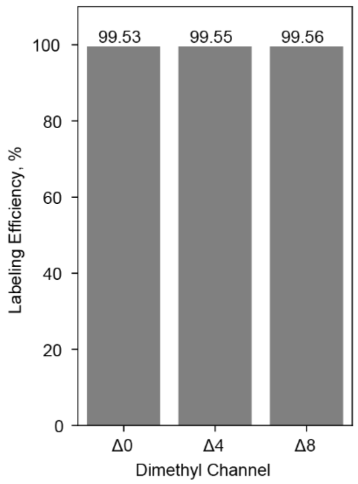


**Fig EV2: Labeling efficiency of tryptic HeLa peptides acquired on the Orbitrap platform.** Tryptic peptides from HeLa cells were labeled with dimethyl mass tags Δ0, Δ4 and Δ8 and acquired individually in DDA mode. Labeling efficiencies based on intensity ratios of labeled peptides relative to all detected peptides are shown.

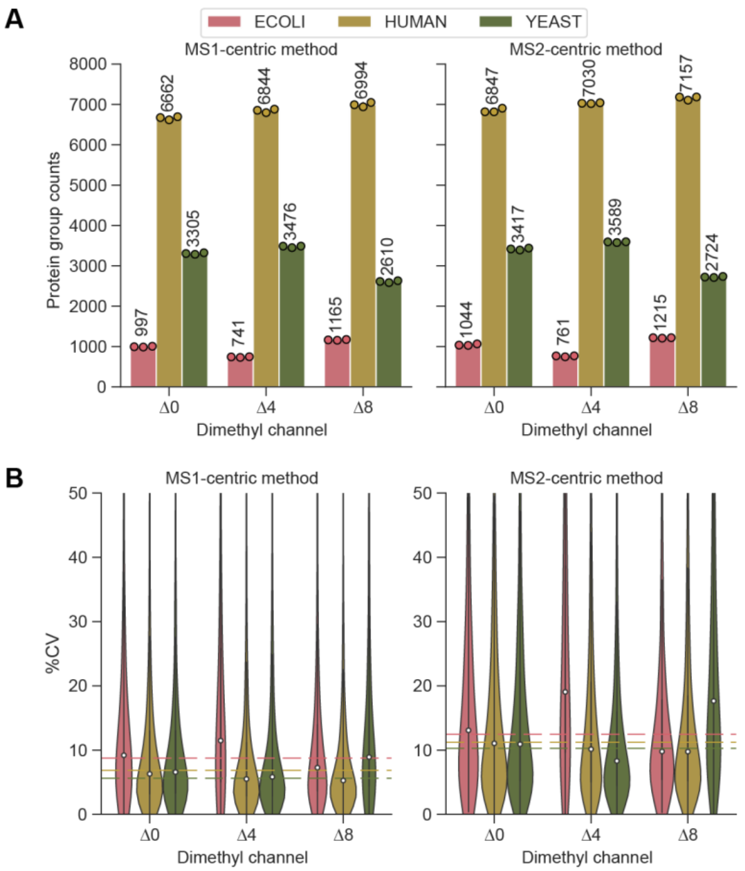


**Fig EV3: Identification rates and coefficient of variation (CVs) in the mixed species experiment.**

A. Tryptic peptides from E. coli, yeast and human were combined at defined ratios before dimethyl labeling. Side-by-side comparison of the number of quantified protein groups for the individual species with MS1- and the MS2-centric methods. 100 ng of peptides were injected per channel.

B. Coefficients of variation (CVs, %) of protein groups shown in (A). Median CVs are shown as dashed lines.

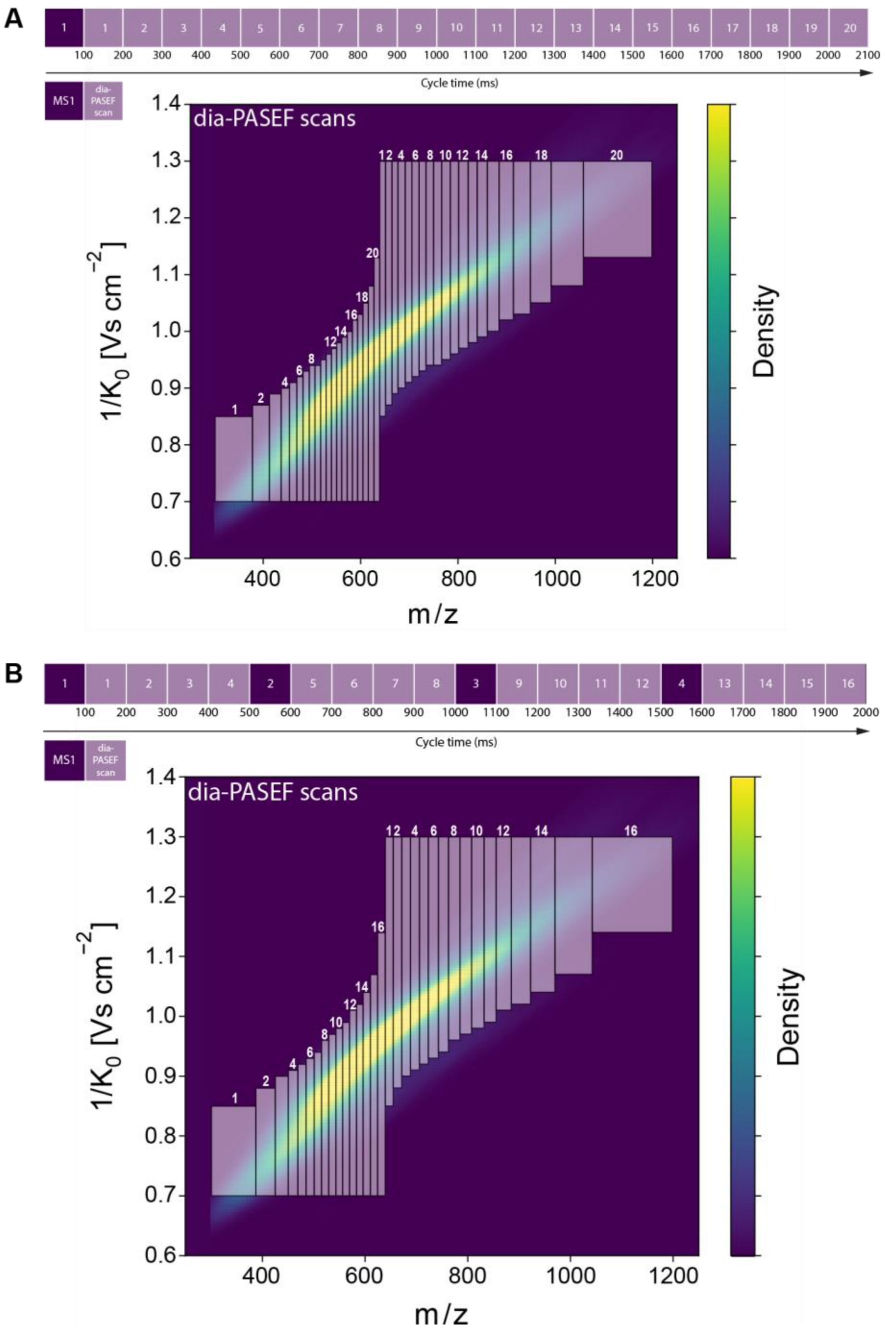


**Fig EV4: dia-PASEF acquisition methods optimized for dimethylated tryptic HeLa peptides.**
A. 20 dia-PASEF scan method used for MS2-centric acquisitions. It consists of one MS1 scan followed by 20 dia-PASEF scans with variable m/z isolation widths and two ion mobility windows in one cycle (~2.1 seconds).
B. 16 dia-PASEF scan method used for MS1-centric acquisitions. This method consists of 16 dia-PASEF and four MS1 scans, with each MS1 scan followed by 4 dia-PASEF scans. Similar to the 20 dia-PASEF scan method (A), the cycle time is about two seconds.

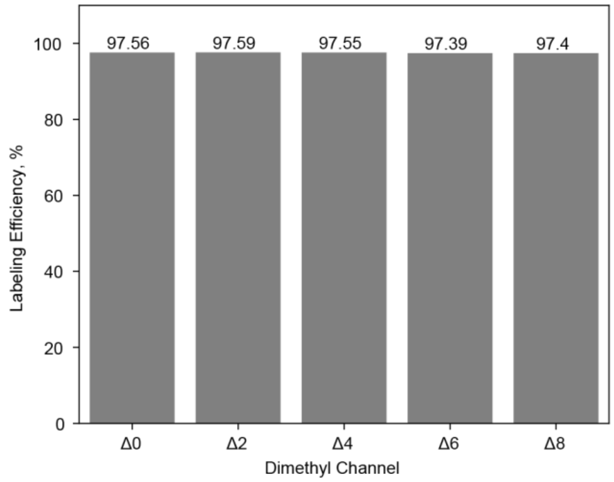


**Fig EV5: Labeling efficiency of Lys-N-derived HeLa peptides acquired on the Orbitrap platform.** Lys-N-derived HeLa peptides from HeLa cells were labeled with dimethyl mass tags Δ0, Δ2, Δ4, Δ6, and Δ8 and acquired individually in DDA mode. Labeling efficiencies based on intensity ratios of labeled peptides relative to all detected peptides are plotted.

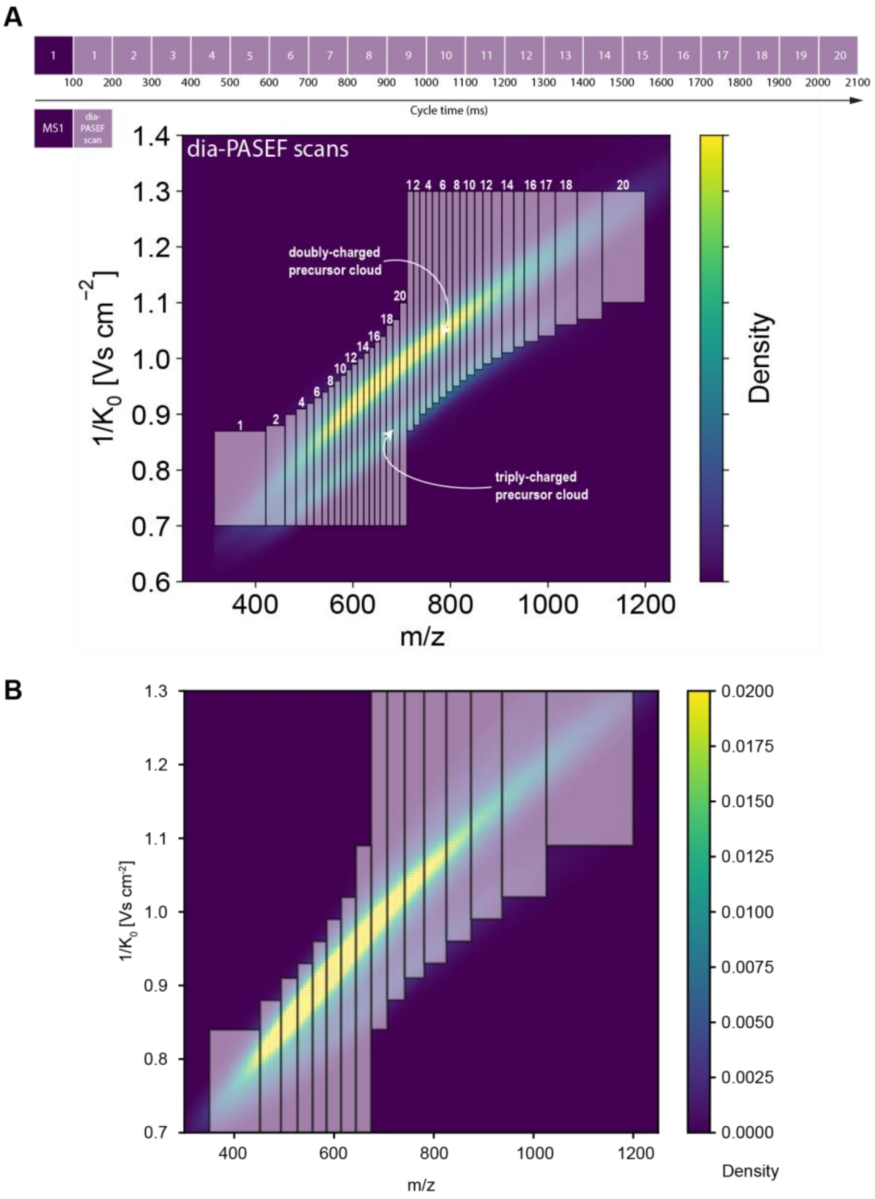


**Fig EV6: Optimal dia-PASEF acquisition method for Lys-N-derived HeLa peptides and tryptic HeLa for ultra-high sensitivity measurements.**

A. 20 dia-PASEF scan method was used for the acquisition of Lys-N-derived HeLa peptides. The method was optimized to specifically cover both doubly- and triply-charged precursors. One cycle time is about 2.1 seconds.

B. Eight dia-PASEF scan method for ultra-high sensitivity measurement of single-cell equivalents and single cells using the 'Whisper40 SPD' gradient in combination with an Aurora Elite LC column (IonOpticks). The acquisition scheme is plotted on top of a kernel density distribution of the precursors from a representative tryptic 48 high-pH fraction HeLa library (Skowronek *et al*, 2022).

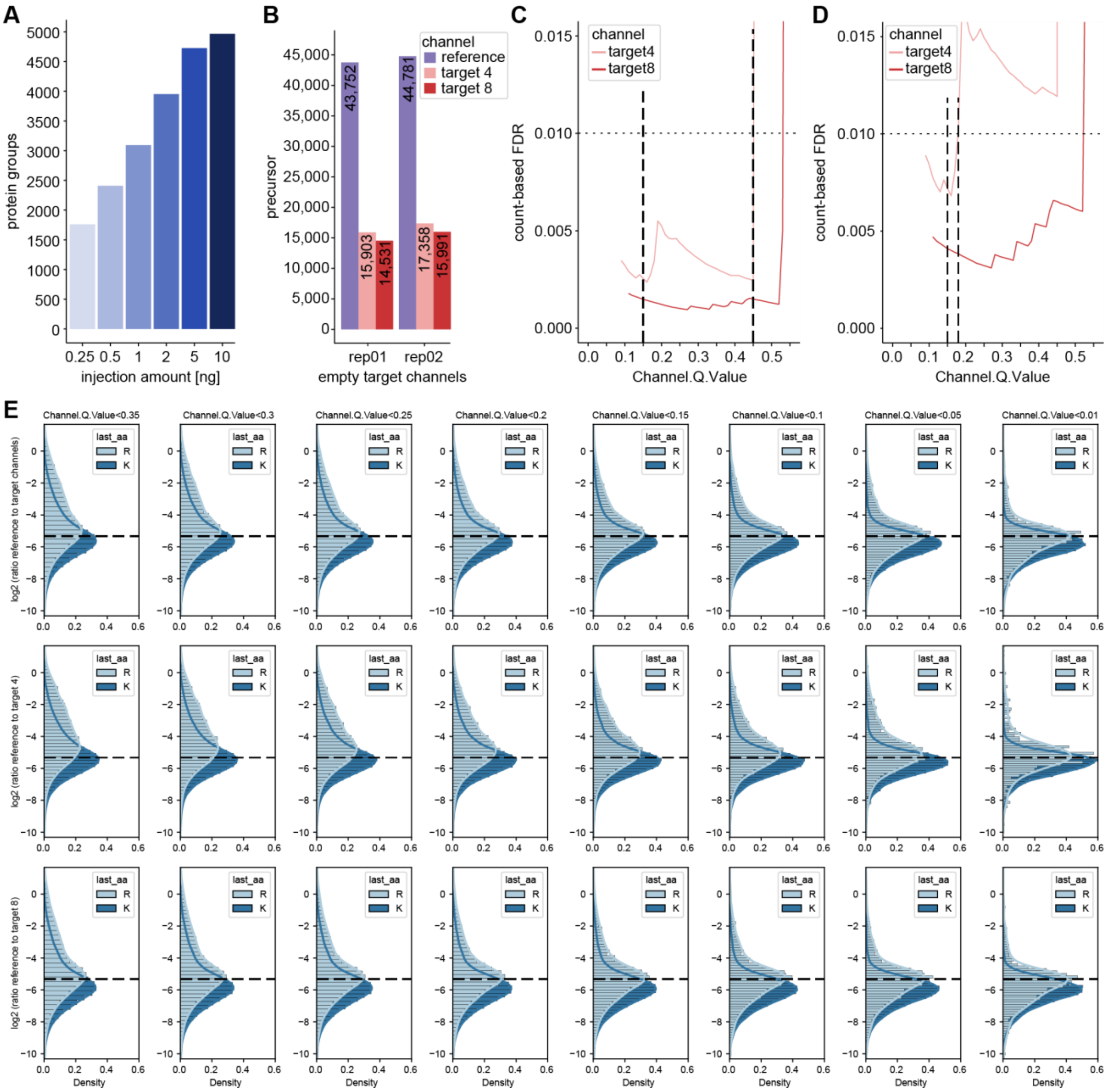


**Figure EV7: Evaluation of 'Translated.Q.Value' and 'Channel.Q.Value' on identifications in empty target channels and quantification in single-cell equivalents.**

A. Dilution series of HeLa peptides to define protein identifications in mDIA workflow, in which the linear increase with input amount levels off at about 10 ng.

B. Precursor identifications of empty target channels at 1% 'Translated.Q.Value' revealed more than 1% false positives.

C and D. Count-based FDR estimation on precursor (C) and protein level (D) by dividing two quantities, namely the maximum number of identified precursors or proteins in four empty target channel runs by the minimum number of identified precursors or proteins in a total of four target channels (target channel 4 and 8) which contain single-cell equivalent amounts. The count-based FDR reveals a ‘Channel.Q.Value’ filter of 0.45 for precursors and 0.17 for proteins at 1%. E. Quantification evaluation of ‘Channel.Q.Value’ filter. Ratios of reference to target channel (top), channel 4 (middle) and 8 (bottom) by RefQuant (see Fig 5) are shown for arginine and lysine precursor at a given filter of ‘Channel.Q.Value’. At all filtering steps, the modes of the distributions are close to the expected ratio (dashed line) with a systematic skew towards lower ratios, which is more pronounced for the arginine precursors. The skews in the distributions are decreased with increased filtering stringency in a gradual manner. The mean quantification ratios are as expected, whereas they start to deviate at 0.2.

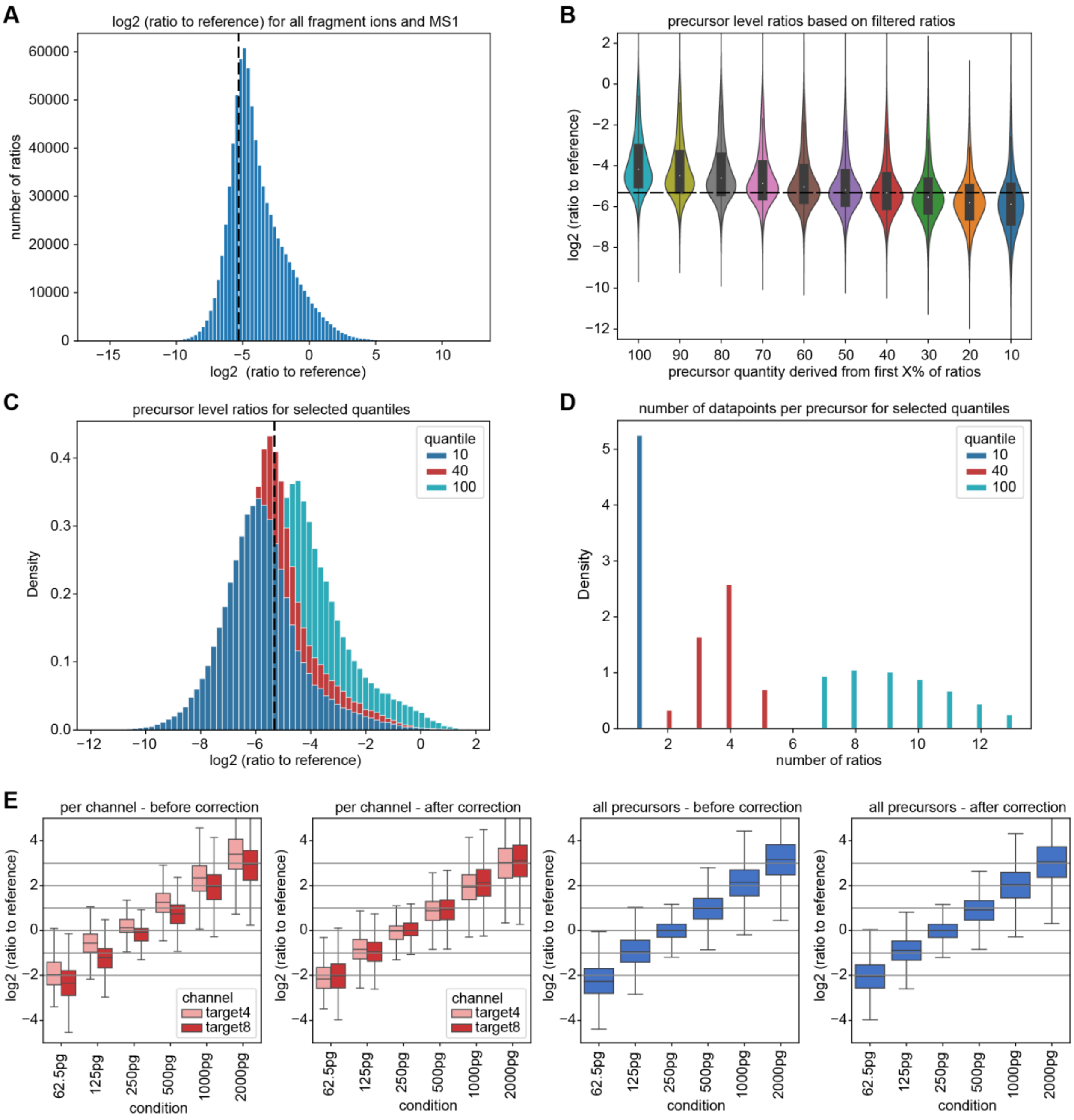


**Figure EV8: Ratios to reference channel evaluation and channel correction comparison in scQuant.**

A. Ratios to the reference for all available ions (fragment ions and MS1 peaks), revealing a distribution that has a mode proximate to the expected ground truth (dashed line). The distribution is asymmetric with a skew towards less extreme ratios. A potential explanation for this skew is that noise or interferences are dominant for a fraction of the ratios (a ratio of 0 might be a 'noise vs. noise' comparison).

B. An approach to mitigate this asymmetry is to filter out some ions before estimating the precursor level ratio. For this, the ratios of a precursor are sorted ascending and only the first ratios are retained (up to the ‘X value’ indicated). The precursor ratio is estimated by taking the mean of the remaining ratios. We see that retaining 40% of the ratios matches the ground truth well and is more symmetric.

C. A more detailed comparison of the distributions when taking 10% or 40% of ratios as compared to all ratios.

D. Number of available ratios per precursor after filtering. We see that in general between 7 to 14 ratios are available, which reduces to 2 to 5 when taking the 40% quantile.

E. Channel correction comparison in scQuant dataset in comparison of all precursor and target channel. Ratio of reference channel to target channel was calculated using RefQuant. All log2 ratios were normalized to the single-cell equivalent sample in the target channel (250 pg). A basic median normalization between the channels was applied.

**Supplemental Text EV9**

**Fundamental considerations about the noise in target and reference for ratio-based quantification**

We investigated to what extent, if any, the ratio between reference and target channels would contribute to variance in the experimental values. If a given peptide had a 50% error in its estimated signal in a 40:1 reference to target ratio, this would manifest in a 20.5-fold ratio in a first run. In a second run – given that MS signals are very reproducible – we would then expect a ratio of 20 or perhaps 19.2. From this it follows that - concerning the noise - the variance of the ratio estimation over runs is not determined by magnitude of the ratio. Following this reasoning, the overall variance of the log2 ratios ${\sigma^2}_{ratio}$ is ${\sigma^2}_{ratio} = {\sigma^2}_{target} + {\sigma^2}_{reference}$ , with ${\sigma^2}_{target}$ denoting the variance of the target channel and ${\sigma^2}_{reference}$ denoting the variance of the reference channel. As the higher abundances of the reference channel stabilizes its signal, we conclude that the overall variation of the ratios will decrease with higher reference proteome amounts.

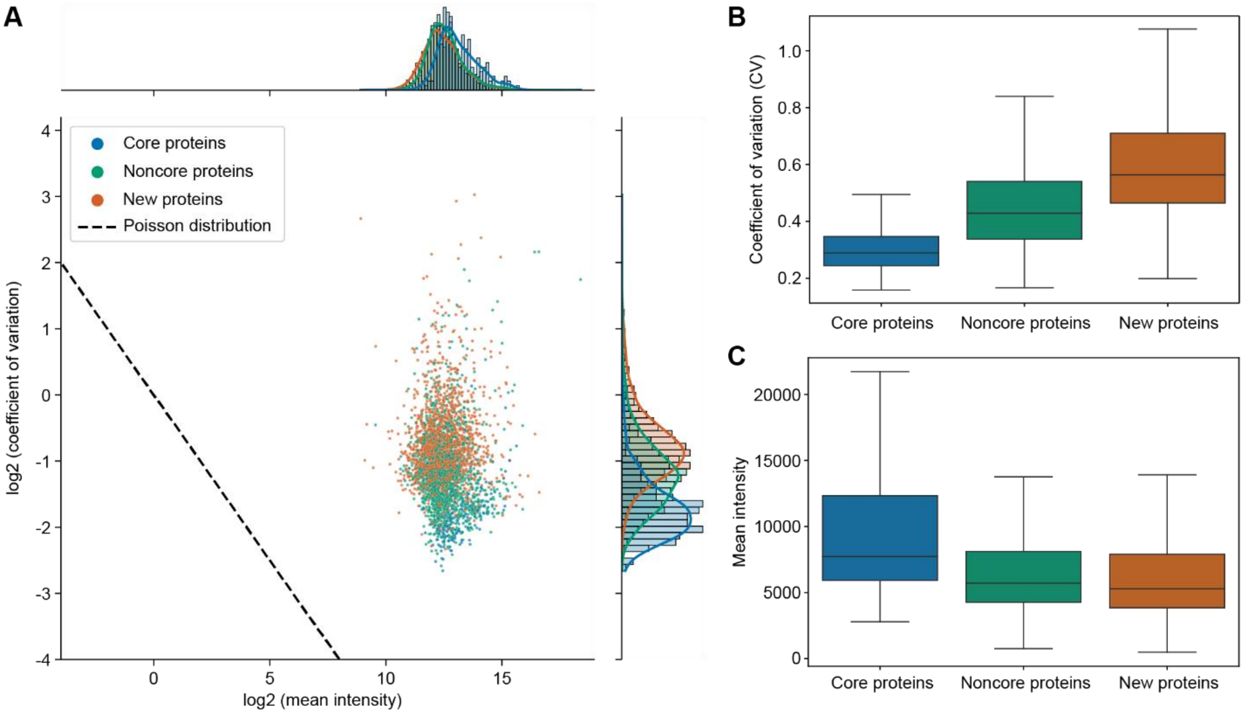


**Figure EV10: mDIA shows that the concept of a stable proteome is still valid at higher proteomic depth in single cells.**

A. Coefficients of variation of single-cell mDIA protein expression plotted against the mean intensity of each protein. ‘Core proteins’ are labeled in blue, ‘noncore proteins’ in green and additionally identified and quantified proteins compared to our previous single-cell publication in orange (Brunner *et al*, 2022).

B. Distribution of coefficients of variation across the mDIA single-cell dataset for ‘core’ (blue), ‘noncore’ (green) and ‘new’ (orange) proteins.

C. Distribution of the mean MS intensity across the mDIA single-cell dataset for ‘core’ (blue), ‘noncore’ (green) and ‘new’ (orange) proteins.
